# From Screening to Differential Diagnosis: Benchmarking 3D Facial Representations Across Craniofacial Conditions

**DOI:** 10.64898/2026.11.12.766227

**Authors:** Tingting Chen, Mian Umair Ahsan, Justin Cotney, Eric C. Liao, Kai Wang

## Abstract

Rare genetic disorders often present with highly heterogeneous physical features: some have distinctive craniofacial characteristics, whereas others produce subtler or more variable facial differences, making phenotype-based diagnosis challenging. Three-dimensional (3D) facial scanning is a noninvasive approach that captures facial shape and surface geometry, providing quantitative information on craniofacial morphology beyond conventional 2D photographs. Here, we investigated how well 3D facial representations support clinical tasks ranging from broad screening to fine-grained differential diagnosis without landmark registration, dense correspondence, or template fitting. We benchmarked four 3D models, including PointNet, PointNet++, DGCNN, and a geometric baseline, using 26,990 facial scans from 18,785 subjects across four prediction tasks. Across models, performance declined as greater diagnostic specificity was required. Broad affected-versus-unaffected screening achieved high performance (accuracy 0.893; AUC 0.956), although sensitivity analyses revealed substantial influences of cohort composition and demographic differences. Fine-grained syndrome diagnosis remained more challenging, with limited top-1 accuracy despite meaningful classwise discrimination. Combining repeated scans improved patient-level diagnosis beyond differences in patient weighting. Prototype inference further improved diagnostic prioritization, placing the correct syndrome among the top five candidates for about 65% of held-out patients across 49 diagnoses. Patient-to-patient retrieval identified phenotypically similar confirmed cases, while useful syndrome prototypes could be constructed from relatively few reference patients. Anatomical region-occlusion analysis revealed syndrome-dependent patterns of facial importance broadly consistent with clinically recognized craniofacial features, although overlapping facial phenotypes contributed to diagnostic confusion. Overall, these findings support the potential of 3D facial representations for diagnostic candidate prioritization and retrieval, while emphasizing the influence of cohort composition on screening performance and the challenges of distinguishing phenotypically similar genetic syndromes.

## 1 Introduction

Craniofacial morphology is an important clinical clue across a broad range of congenital and genetic conditions [1, 2]. Orofacial clefts produce readily observable structural abnormalities [3, 4], whereas many rare genetic syndromes are associated with subtler and more variable combinations of facial features that can help narrow a diagnosis but may require specialized dysmorphology expertise to recognize [2, 5]. Two-dimensional (2D) facial photographs have already shown substantial value for facilitating rare disease diagnosis by enabling quantitative facial phenotyping and prioritization of candidate syndromes [5–10]. However, conventional frontal photographs capture only a limited projection of facial anatomy and do not directly represent three-dimensional (3D) features such as facial depth, contour, asymmetry, and surface geometry [11]. 3D facial scanning can capture this additional structural information, potentially providing a richer representation of craniofacial phenotype [11–13]. In clinical practice, facial phenotyping is generally most useful for identifying atypical morphology and prioritizing conditions for further clinical or genetic evaluation rather than establishing a diagnosis from facial appearance alone [5, 8]. 3D facial scanning is particularly well suited to this setting because it enables rapid, non-ionizing, and quantitative capture of facial surface geometry and is already used in craniofacial and orthodontic practice [14–16].

Building on the richer geometric information captured by 3D facial imaging, computational studies have shown that facial morphology contains information useful for distinguishing craniofacial conditions and genetic syndromes [11, 13, 17–19], complementing advances in image-based facial phenotyping [7, 9]. Most established 3D methods first establish anatomical correspondence using landmarks, dense registration, or statistical shape models [20–22]. Such correspondence enables point-wise comparison of anatomy but also introduces additional preprocessing and dependence on landmark detection or template fitting, which may be particularly challenging for markedly dysmorphic faces with substantial structural abnormalities [23]. Moreover, template fitting may reduce measured deviations from normative facial shape (Supplementary Fig. S1), while approaches that impose facial symmetry or analyze only the symmetric component may discard clinically relevant asymmetry [24]. These limitations motivate alternative approaches that preserve native facial morphology. 3D point-cloud neural networks offer such an alternative by learning directly from facial surface geometry without requiring dense correspondence [25–27]. However, it remains unclear how well these representations support clinically relevant tasks across different levels of diagnostic specificity.

A central question is how these representations behave as the diagnostic task becomes more specific (Fig. 1a). Detecting whether a face shows craniofacial abnormalities is fundamentally different from distinguishing among cleft subtypes or identifying one diagnosis among many rare disorders. Fine-grained syndrome recognition is commonly formulated as closed-set classification, in which the model must assign a single label from a predefined set [7, 13]. In clinical practice, however, facial phenotyping is often used more flexibly: facial morphology can help determine which confirmed diagnostic groups or previously diagnosed patients an individual resembles and which conditions warrant further consideration [8, 9]. A representation may therefore remain clinically informative even when the correct diagnosis is not consistently assigned the highest score, provided that it preserves meaningful discrimination and similarity among candidate conditions [7, 9]. Evaluating diagnostic candidate ranking, retrieval of similar confirmed patients, and the influence of reference-library size may therefore help establish how 3D facial representations could support differential diagnosis beyond conventional single-label classification.

**Figure 1:**
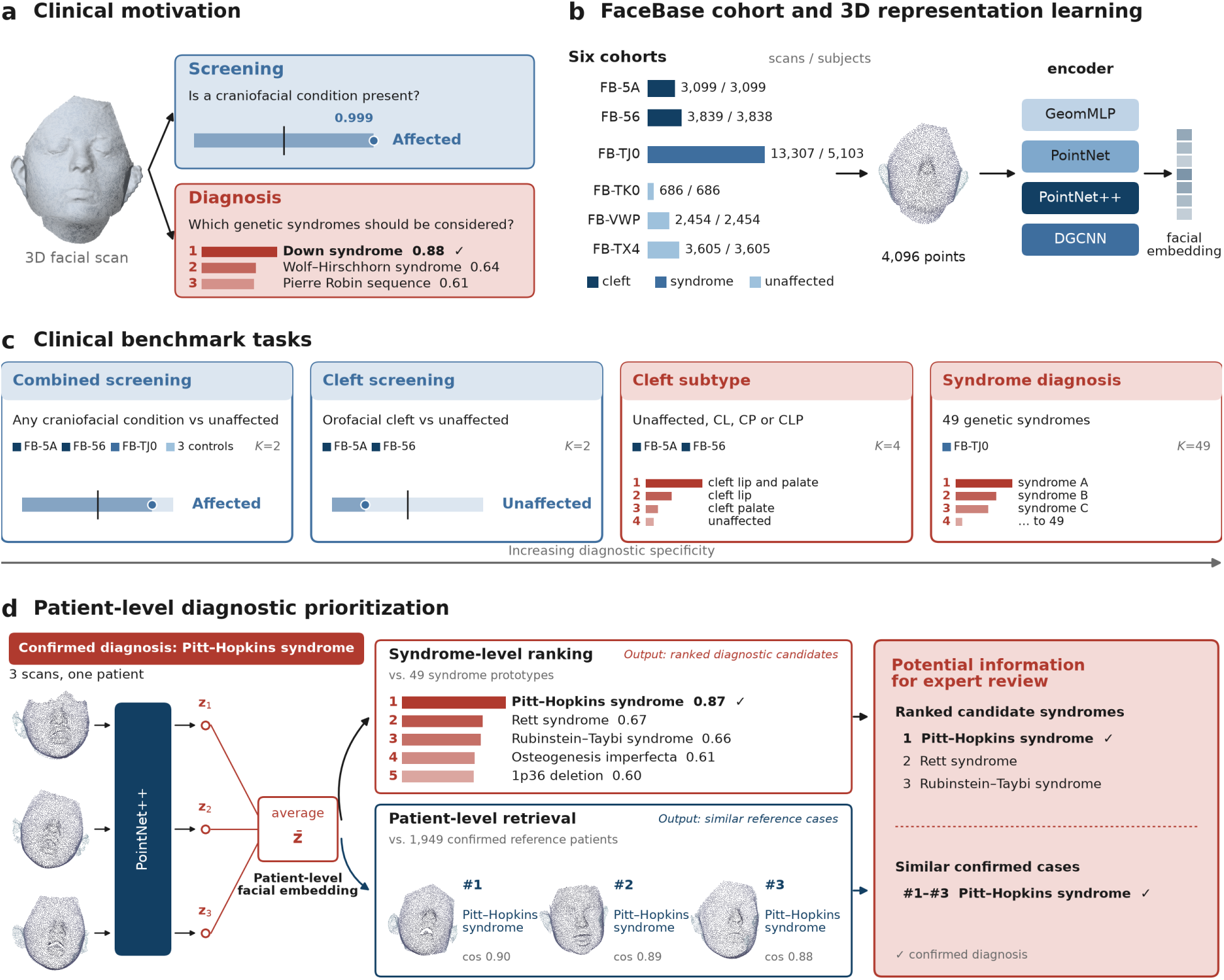
From 3D facial screening to diagnostic prioritization. **(a)** Clinical motivation: screening for craniofacial conditions and prioritizing candidate genetic syndromes. Both examples show PointNet++ predictions for the same held-out FB-TJ0 scan from a patient with Down syndrome. Screening outputs are shown in blue and diagnostic rankings in red; the two tasks were evaluated separately. **(b)** FaceBase cohorts and 3D representation learning. Numbers indicate processed scans and distinct subjects across six cohorts. Each scan was represented by a 4,096-point cloud without facial landmarks, dense correspondence, or template registration. Four encoders (GeomMLP, PointNet, PointNet++, and DGCNN) were evaluated independently. Task-specific subsets were defined according to eligibility criteria (Table 1). **(c)** Four benchmark tasks: combined affected-versus-unaffected screening, cleft screening, four-class cleft subtype classification, and 49-class syndrome diagnosis. The cohorts, numbers of scans and subjects, and number of classes (*K*) are shown for each task. CL, cleft lip; CP, cleft palate; CLP, cleft lip and palate. Score bars and candidate names are schematic. **(d)** Patient-level diagnostic prioritization for a held-out patient with Pitt–Hopkins syndrome and three facial scans, using actual outputs from one trained PointNet++ model. Averaged scan embeddings were compared by cosine similarity with syndrome prototypes and individual training-patient embeddings. The confirmed syndrome ranked first among 49 candidates, and all three nearest reference patients shared the diagnosis. Syndrome-level ranking and patient-level retrieval provide complementary information for potential expert review and were not computationally combined. Quantitative retrieval and ranking results are shown in Fig. 5.

**Table 1:** FaceBase cohort composition and contributions to the clinical benchmark tasks. The benchmark comprised 26,990 processed 3D facial scans from 18,785 subjects across six FaceBase datasets. FB-5A and FB-56 contributed cleft cases, FB-TJ0 contributed individuals with genetic syndromes, and FB-TK0, FB-VWP, and FB-TX4 contributed unaffected controls. Tasks included four-class cleft subtyping (c4), binary cleft screening (c2), 49-class syndrome diagnosis (s49), and combined affected-versus-unaffected screening (b). Counts shown in the table reflect the processed cohort before task-specific eligibility criteria were applied.

| Dataset | Scans | Subjects | Platform | Cohort | Tasks |
|---|---|---|---|---|---|
| FB-5A [29] | 3,099 | 3,099 | 3dMD | Cleft cohort | c4, c2, b |
| FB-56 [30] | 3,839 | 3,838 | 3dMD | Cleft cohort | c4, c2, b |
| FB-TJ0 [31] | 13,307 | 5,103 | 3dMD | Genetic syndromes | s49, b |
| FB-TK0 [32] | 686 | 686 | Unspecified | Unaffected controls | b |
| FB-VWP [33] | 2,454 | 2,454 | Unspecified | Normal facial variation | b |
| FB-TX4 [34] | 3,605 | 3,605 | Unspecified | Unaffected controls | b |
| Total | 26,990 | 18,785 |  |  |  |

A clinically meaningful benchmark must also account for characteristics of craniofacial datasets that can affect measured performance and model interpretation. Differences in cohort composition, age, and other demographic characteristics may influence screening performance, making it important to distinguish facial morphology associated with clinical conditions from cohort-related differences. In addition, individual patients may contribute multiple scans, so scan-level train– test splitting can place different acquisitions from the same person in both partitions and inflate apparent generalization. Conversely, repeated scans may provide complementary observations of the same patient’s morphology when properly combined at inference. Beyond predictive performance, understanding which facial regions contribute to learned representations is important for assessing their clinical relevance. Determining whether models rely on anatomically meaningful features associated with known craniofacial phenotypes can provide insight into their predictions and limitations, especially when different syndromes share similar facial characteristics. Thus, cohort-aware evaluation, subject-independent data partitioning, controlled patient-level aggregation, and anatomical interpretability analysis are complementary components of a clinically meaningful assessment of 3D facial representations.

Here, we systematically evaluate 3D facial representations without template registration across four tasks spanning broad craniofacial screening, cleft screening and subtyping, and fine-grained syndrome diagnosis (Fig. 1). Using 26,990 scans from 18,785 subjects across six FaceBase datasets [28], we compare PointNet [25], PointNet++ [26], DGCNN [27], and a geometric baseline (GeomMLP) under a common preprocessing framework without landmark registration, dense correspondence, or template fitting. We investigate the influence of cohort composition and demographic differences on screening performance and evaluate whether combining repeated scans improves patient-level diagnosis beyond differences in patient weighting. We further assess prototype-based diagnostic candidate ranking, retrieval of similar confirmed patients, and the effect of reference-library size. Anatomical region-occlusion analysis is used to identify facial regions contributing to syndrome-associated representations. Across models, performance declines as greater diagnostic specificity is required, while cohort and demographic differences substantially influence broad screening performance. Combining repeated scans improves fine-grained diagnosis, and prototype inference supports prioritization of candidate syndromes and retrieval of similar confirmed patients. Regional importance patterns broadly align with clinically recognized craniofacial features, although distinguishing syndromes with overlapping facial characteristics remains challenging. These findings highlight the potential of 3D facial representations to complement expert clinical and genetic evaluation through morphological screening and diagnostic prioritization.

## 2 Results

### 2.1 Study cohorts and clinical benchmark tasks

The benchmark included 26,990 3D facial scans from 18,785 subjects across six FaceBase datasets, comprising individuals with orofacial clefts, genetic syndromes, and unaffected controls (Fig. 1b; Table 1). FB-5A [29] and FB-56 [30] provided cleft phenotypes from family-based studies, whereas FB-TJ0 [31] provided diagnosis-level labels for individuals with genetic syndromes. Three additional cohorts (FB-TK0 [32], FB-VWP [33], and FB-TX4 [34]) contributed unaffected individuals to the screening tasks.

The dataset contained substantial repeated imaging: individual subjects contributed as many as 34 scans, with FB-TJ0 alone containing 13,307 scans from 5,103 subjects. To prevent information leakage between training and evaluation, data were partitioned at the subject level, ensuring that scans from the same individual did not appear in different partitions. We also examined whether combining repeated scans improved patient-level diagnostic performance.

We evaluated four tasks spanning increasing levels of diagnostic specificity: combined affected-versus-unaffected screening, binary cleft screening, four-class cleft subtype classification, and 49-class genetic syndrome diagnosis (Fig. 1c). Task-specific eligibility criteria resulted in different subsets of the processed cohorts. All models operated directly on sampled 3D facial points without landmark registration, dense anatomical correspondence, or normative template fitting.

### 2.2 Diagnostic performance declines with increasing task specificity

We compared four approaches for representing 3D facial shape: a multilayer perceptron applied to 26 handcrafted global geometric descriptors (GeomMLP), PointNet [25], PointNet++ [26], and DGCNN [27]—across the four clinical benchmark tasks.

Performance varied substantially more across clinical tasks than across model architectures (Fig. 2). On the combined affected-versus-unaffected screening task, PointNet++ achieved an accuracy of 0.893 *±* 0.007 and an AUC of 0.956 *±* 0.002. Binary cleft screening also showed relatively high performance, with PointNet++ achieving 0.872 *±* 0.015 accuracy and 0.914 *±* 0.012 AUC. DGCNN reached a similar AUC but lower accuracy (0.815 *±* 0.018 accuracy; 0.914 *±* 0.019 AUC), whereas GeomMLP achieved a lower AUC of 0.726 *±* 0.002. These results indicate that broad differences in craniofacial morphology were detectable across independent training runs, although combined screening performance requires cautious interpretation because of cohort composition and demographic differences (Section 2.3).

**Figure 2:**
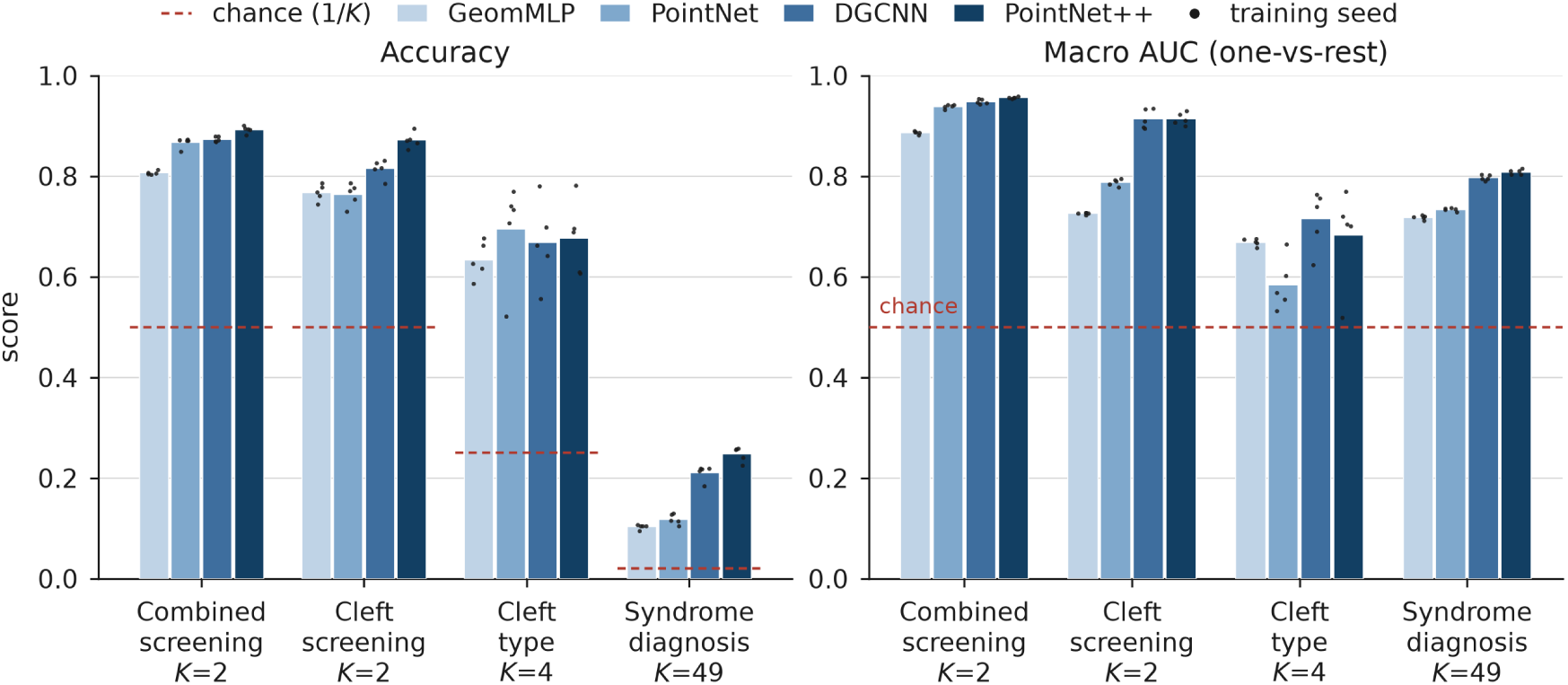
Performance across four craniofacial analysis tasks and four 3D facial representations. Scan-level accuracy (left) and macro one-versus-rest AUC (right) are shown for GeomMLP, PointNet, DGCNN, and PointNet++ using softmax inference. Bars show the mean across five independently trained models with different random seeds, and dots indicate individual training runs. The same subject-level data partitions were used for all seeds. Chance accuracy is 1*/K*, where *K* is the number of classes, and chance AUC is 0.5. Performance decreased as the task progressed from binary screening to fine-grained syndrome diagnosis, whereas macro AUC remained comparatively high on the 49-class task. This pattern was consistent across architectures and independent training runs.

Performance declined as greater diagnostic specificity was required. Within the cleft cohorts, four-class subtyping was more difficult than binary cleft screening, with PointNet++ macro AUC decreasing from 0.914 *±* 0.012 to 0.683 *±* 0.096. The 49-class syndrome-diagnosis task was more challenging, with PointNet++ achieving a scan-level top-1 accuracy of 0.248 *±* 0.015. This progression was observed across architectures, with PointNet++ and DGCNN generally performing best among the learned models. Differences between architectures were modest relative to the performance changes across clinical tasks.

Despite limited top-1 accuracy, the syndrome-diagnosis models retained substantial class-wise discrimination. PointNet++ achieved a macro one-versus-rest AUC of 0.808 *±* 0.005 on the 49-class task (Table 2; Fig. 2), a pattern observed across all five training runs and multiple architectures. Moreover, the correct syndrome appeared among the five highest-ranked candidates for 0.540 *±* 0.012 of scans. Thus, although distinguishing one syndrome from many competing diagnoses remained difficult, the models retained information that could support diagnostic candidate prioritization. Overall, these findings highlight the substantial difference in performance between broad cranio-facial screening and fine-grained syndrome diagnosis. The gap between class-wise discrimination and top-1 accuracy suggests that learned facial representations retain diagnostic information even when the correct diagnosis is not ranked first, motivating following further analyses to improve diagnostic prioritization.

**Table 2:**
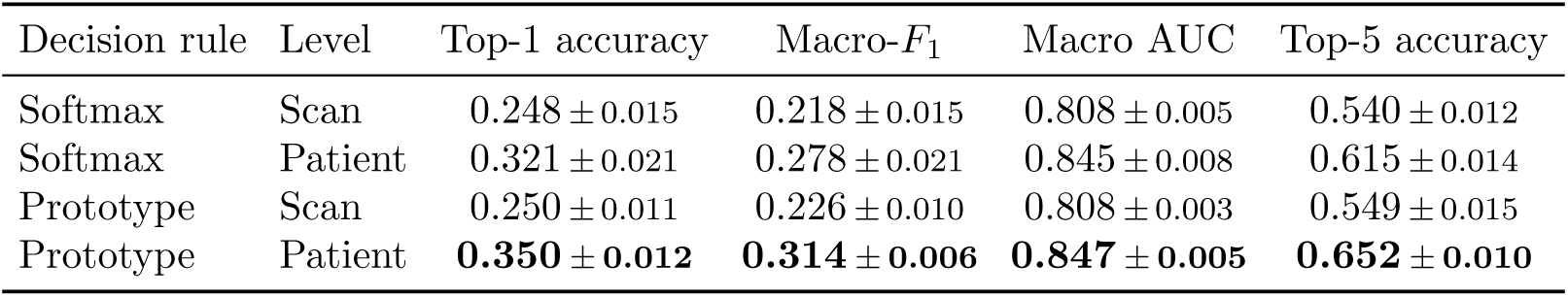
Effect of patient-level aggregation and inference strategy on fine-grained diagnosis. Performance is shown for the 49-class syndrome-diagnosis task using PointNet++ representations. Values are mean *±* standard deviation across five independently trained models. Scan-level predictions treat each acquisition independently, whereas patient-level predictions combine repeated scans from the same individual. For softmax inference, class probabilities are averaged across scans; for prototype inference, scan embeddings are averaged before comparison with class prototypes. Top-1 accuracy is the proportion of scans or subjects whose highest-scoring predicted diagnosis is correct, while top-5 accuracy is the proportion for which the true diagnosis appears among the five highest-scoring classes (chance = 5*/*49 = 0.102). Macro-*F*_1_ and macro AUC average class-wise *F*_1_ scores and one-versus-rest ROC AUC values, respectively, giving equal weight to each diagnosis. Evaluation included 1056 test scans from 385 test subjects.

### 2.3 Cohort composition and demographic differences influence screening performance

The combined screening task pooled six FaceBase cohorts with markedly different diagnostic compositions and demographics (Table 3). Three cohorts (FB-TK0, FB-VWP, and FB-TX4; 6,459 subjects) contributed only unaffected individuals, whereas 77.7% of FB-TJ0 subjects were affected, compared with 22.0% in FB-5A and 14.6% in FB-56. Source cohort was therefore strongly associated with diagnostic status. A baseline using only source-cohort membership, without facial information, achieved an AUC of 0.912, compared with 0.954 *±* 0.004 for PointNet++ on the same held-out subjects. This indicates that cohort composition was strongly associated with diagnostic status and could substantially inflate apparent screening performance. In addition, affected individuals were substantially younger than unaffected individuals within all three cohorts containing affected cases (median age 8–12 versus 28–36 years; Fig. 3a).

**Figure 3:**
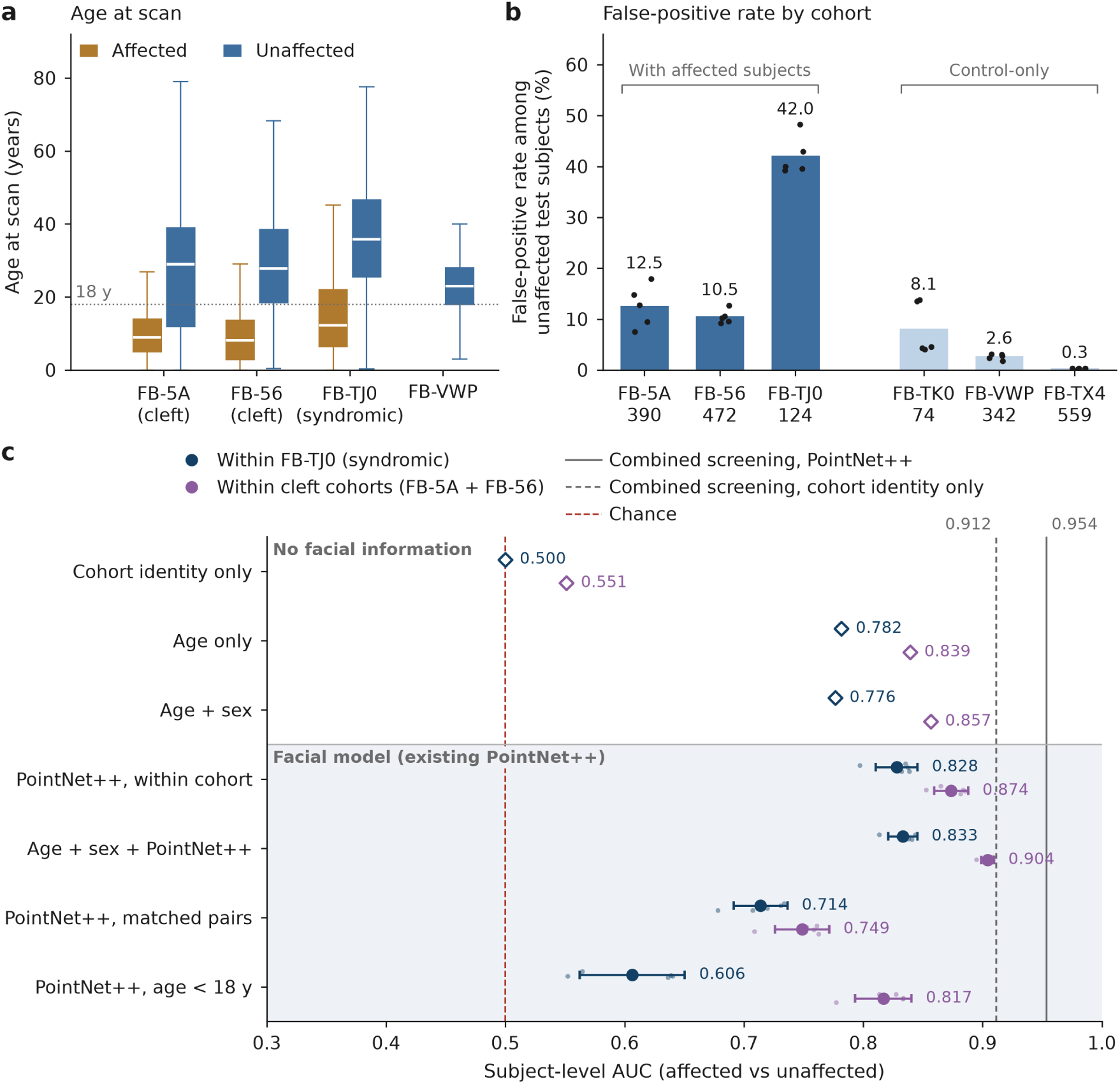
Cohort composition and demographic differences influence combined affected-versus-unaffected screening. All analyses used the existing combined-task PointNet++ models and subject-level test-set predictions; no model was retrained, and only the subset of test subjects or covariates entering the analysis changed. (a) Age at scan by source cohort and diagnostic status for cohorts with available age metadata. Boxes show the interquartile range, center lines the median, and whiskers 1.5*×* the interquartile range. (b) False-positive rate among unaffected test subjects at the default decision threshold. Bars show the mean across five trained models and dots show individual models; *n* indicates the number of unaffected test subjects in each cohort. (c) Subject-level AUCs for affected-versus-unaffected discrimination under increasingly controlled comparisons. Open diamonds denote predictors without facial information (cohort identity, age alone, and age+sex), whereas filled circles and horizontal bars denote mean *±* SD across the five PointNet++ models or analyses incorporating the existing PointNet++ score. Vertical dashed lines indicate chance (0.5), the cohort-identity-only baseline, and the combined all-cohort PointNet++ AUC.

**Table 3:** Composition of the combined affected-versus-unaffected screening task by source cohort. Subject and scan counts correspond to the combined screening task, a subset of the processed corpus in Table 1. Test subjects are those assigned to the held-out test partition. Age is reported in years as median [interquartile range], and demographic percentages are calculated within each cohort and diagnostic group. Of the 978 unaffected FB-TJ0 subjects, 821 were relatives of affected individuals and 157 were unrelated controls. Age and sex metadata were unavailable for FB-TK0 and FB-TX4. Dashes indicate unavailable information.

| Cohort | Diagnostic status | Subjects | Scans | Test subjects | Age, y | <18 y (%) | Female (%) |
|---|---|---|---|---|---|---|---|
| FB-5A (OFC1) | Affected | 683 | 683 | 96 | 9.0 [5.0–14.0] | 83.2 | 43.0 |
|  | Unaffected | 2,415 | 2,415 | 390 | 29.0 [12.0–39.0] | 33.1 | 58.9 |
| FB-56 (OFC2) | Affected | 551 | 552 | 77 | 8.1 [3.0–13.6] | 84.6 | 48.3 |
|  | Unaffected | 3,223 | 3,223 | 472 | 27.8 [18.5–38.5] | 23.5 | 57.7 |
| FB-TJ0 (syndromic) | Affected | 3,414 | 9,044 | 524 | 12.2 [6.4–22.0] | 67.7 | 53.8 |
|  | Unaffected | 978 | 2,236 | 124 | 35.8 [25.6–46.7] | 13.0 | 62.7 |
| FB-TK0 | Unaffected | 404 | 404 | 74 | – | – | – |
| FB-VWP (Facial Norms) | Unaffected | 2,452 | 2,452 | 342 | 23.0 [18.0–28.0] | 23.7 | 61.3 |
| FB-TX4 (Tanzania) | Unaffected | 3,603 | 3,603 | 559 | – | – | – |

To reduce the influence of between-cohort differences, we re-evaluated the existing PointNet++ predictions within individual cohorts, without retraining. AUC decreased to 0.828 *±* 0.017 within FB-TJ0 and to 0.874 *±* 0.014 within the combined cleft cohorts (FB-5A, 0.861 *±* 0.025; FB-56, 0.882 *±* 0.009; Fig. 3c; Supplementary Table S1). The same pattern was reflected in false-positive rates at the default threshold: 42.0% of unaffected FB-TJ0 subjects were classified as affected, compared with 12.5% and 10.5% in FB-5A and FB-56, and 8.1%, 2.6%, and 0.3% in FB-TK0, FB-VWP, and FB-TX4 (Fig. 3b). Thus, unaffected subjects recruited alongside affected individuals were more frequently classified as affected than those from unaffected-only cohorts.

Age alone was also predictive of affected status, yielding AUCs of 0.782 within FB-TJ0 and 0.839 within the cleft cohorts (Fig. 3c). In a secondary cross-validated analysis using existing held-out PointNet++ scores, adding the facial-model score to age and sex improved AUC from 0.776 to 0.833 *±* 0.012 within FB-TJ0 and from 0.857 to 0.904 *±* 0.005 within the cleft cohorts. In 1:1 age- and sex-matched comparisons within cohort, the facial score retained an AUC of 0.749 *±* 0.023 among 147 matched cleft pairs and 0.714 *±* 0.023 among 107 matched FB-TJ0 pairs. However, the matched FB-TJ0 comparison was largely restricted to adults because few unaffected children were available. Among FB-TJ0 subjects younger than 18 years, only 22 unaffected individuals were available, and discrimination was weak and imprecise (AUC 0.606 *±* 0.044; 95% CI for the 25-model ensemble, 0.489–0.761), whereas the corresponding AUC in the cleft cohorts was 0.817 *±* 0.024.

Overall, these analyses show that our combined screening AUC is substantially influenced by cohort composition and demographic differences. Although facial representations retained discriminatory information in more restricted comparisons, the high combined-screening AUC should not be interpreted as evidence of generalizable clinical screening performance, particularly for genetic syndromes in children.

### 2.4 Combining repeated scans improves patient-level diagnosis

Because many individuals in the FB-TJ0 syndrome cohort contributed repeated 3D facial scans, we investigated whether combining scans improved fine-grained diagnosis. Patient-level aggregation increased top-1 accuracy by +7.3 and +10.0 percentage points under softmax and prototype inference, respectively, with corresponding improvements in macro-*F*_1_ and macro AUC (Table 2; Fig. 4a). However, scan-level evaluation gives greater weight to patients with more scans, whereas patient-level evaluation counts each patient equally. Thus, these improvements may partly reflect differences in patient weighting rather than aggregation itself. In contrast, aggregation provided little additional benefit for combined affected-versus-unaffected screening (Supplementary Table S2).

**Figure 4:**
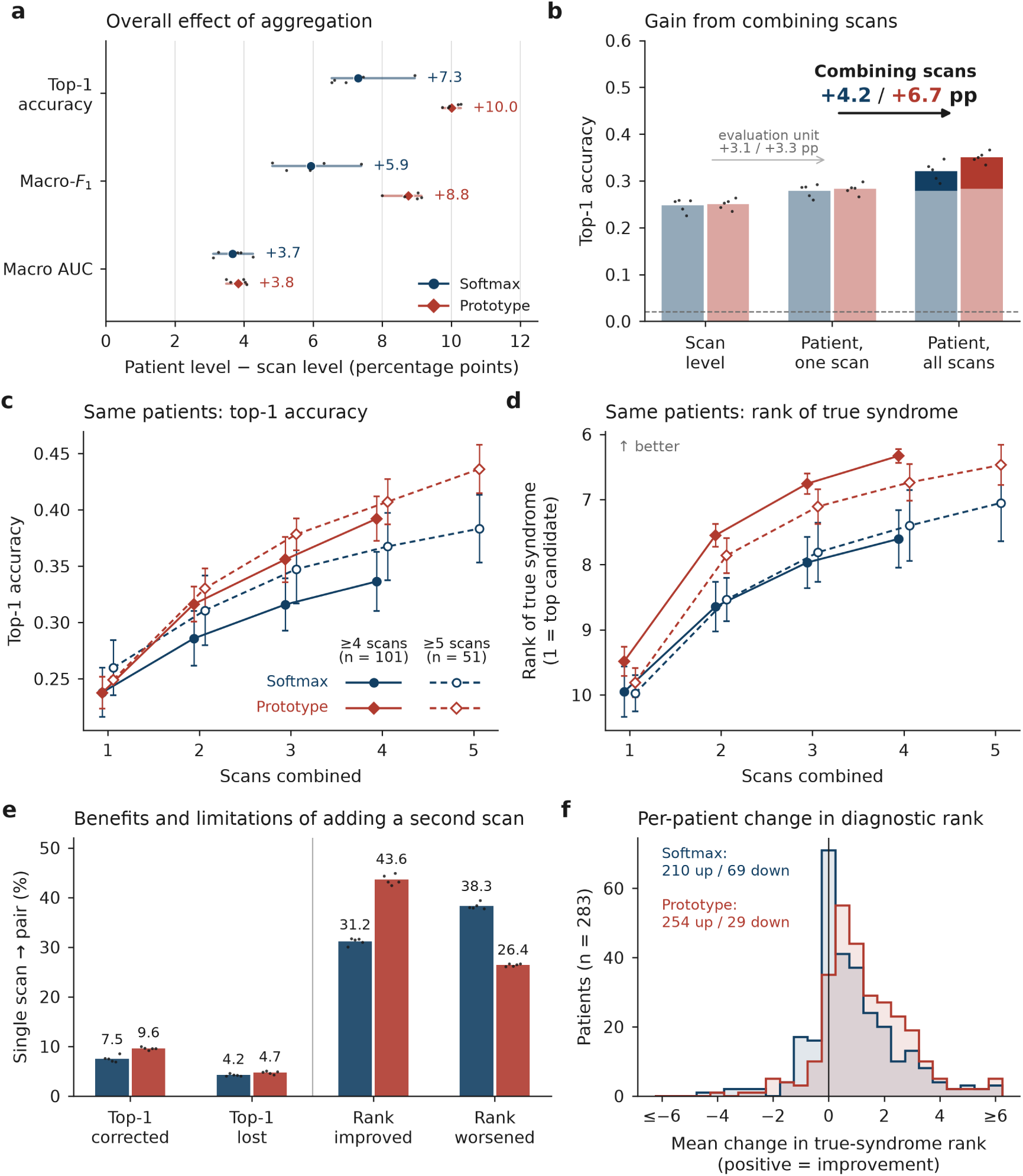
Combining repeated facial scans improves patient-level diagnosis. Results for the 49-class syndrome-diagnosis task using five independently trained PointNet++ models without retraining. Navy indicates softmax inference; red indicates prototype inference. Values are mean *±* SD across five trained models, with inference seeds averaged within each model; dots indicate individual models. **(a)** Overall changes in top-1 accuracy, macro-*F*_1_, and macro AUC from scan-level to patient-level evaluation (385 patients, 1056 scans). Horizontal lines indicate the range across models, and labels show changes in percentage points. **(b)** Decomposition of top-1 accuracy gains into changes in patient weighting (scan-level versus patient-level evaluation using one randomly selected scan) and the additional benefit of combining all available scans. Darker bar segments indicate the gains from combining scans; annotations report changes in percentage points under softmax and prototype inference, respectively. **(c,d)** Top-1 accuracy (c) and mean rank of the true syndrome among 49 candidates (d; rank 1 is best) as a function of the number of scans combined. Solid lines represent patients with at least four scans (*n* = 101), and dashed lines represent those with at least five scans (*n* = 51). Comparisons within each curve use the same patients. **(e)** Changes in top-1 correctness and true-syndrome rank when adding a second scan, expressed as percentages of single-scan-to-pair comparisons with patients weighted equally. **(f)** Distribution of per-patient mean changes in true-syndrome rank after adding a second scan (*n* = 283 patients). Changes were averaged across scan pairs, trained models, and inference seeds; positive values indicate improved ranking. Values outside *±*6 are included in the end bins.

To separate these effects, we evaluated the existing PointNet++ models using one randomly selected scan per patient and then all available scans, without retraining. Changing the evaluation unit alone increased top-1 accuracy by 3.1 and 3.3 percentage points under softmax and prototype inference, respectively. Combining scans provided additional gains of 4.2 and 6.7 percentage points, corresponding to approximately 57% and 67% of the overall patient-level improvements (Fig. 4b). These results demonstrate that combining repeated scans contributes additional diagnostic information beyond differences in how patients are weighted.

We next examined how diagnostic performance changed with increasing scan count using fixed patient cohorts (Fig. 4c,d). Among 101 patients with at least four scans, top-1 accuracy increased from 0.238 to 0.336 under softmax inference and from 0.238 to 0.392 under prototype inference when combining one versus four scans. The mean rank of the true syndrome among 49 candidates also improved under both inference strategies. Across 283 patients with at least two scans, adding a second scan increased top-1 accuracy by 3.3 percentage points (95% CI, 2.4–4.2) under softmax inference and 4.8 percentage points (95% CI, 3.7–6.1) under prototype inference. It also moved the true syndrome higher in the candidate list by 0.86 and 1.30 positions, respectively. Performance generally improved with additional scans, with the largest gains observed early and smaller incremental benefits thereafter. Similar trends were observed among patients with at least five scans (Supplementary Table S3).

The benefit of a second scan varied across patients (Fig. 4e,f). Adding a second scan corrected an incorrect top-1 prediction in 7.5% of comparisons under softmax inference and 9.6% under prototype inference, but also caused previously correct predictions to become incorrect in 4.2% and 4.7%, respectively. Changes in diagnostic ranking were more frequent than changes in top-1 assignment. Under prototype inference, the true syndrome moved higher in 43.6% of comparisons and lower in 26.4%, whereas softmax inference showed more frequent downward than upward movements. Nevertheless, averaging rank changes within each patient showed an overall positive effect for 210 of 283 patients under softmax inference and 254 under prototype inference (Fig. 4f).

These findings indicate that combining repeated facial scans improves patient-level diagnostic prioritization beyond differences in patient weighting. The largest gains were generally observed when combining the first few scans, particularly under prototype inference, although additional scans did not benefit every patient.

### 2.5 Prototype inference supports diagnostic prioritization and patient retrieval

Having established the benefit of patient-level aggregation, we examined whether prototype inference could better support fine-grained diagnostic prioritization than the conventional softmax classifier. At the patient level, prototype inference improved top-1 accuracy from 0.321 to 0.350, macro-*F*_1_ from 0.278 to 0.314, and top-5 accuracy from 0.615 to 0.652, while macro AUC remained largely unchanged (0.845 versus 0.847; Table 2). These gains varied across diagnoses, with per-class *F*_1_ increasing for 31 diagnoses and decreasing for 9 (Supplementary Figs. S2 and S3). Thus, prototype inference modestly improved diagnostic assignment and ranking rather than uniformly improving discrimination across syndromes.

Because facial phenotyping is primarily used to narrow a differential diagnosis rather than establish a diagnosis from facial appearance alone, we further evaluated whether the learned representations could prioritize candidate syndromes for expert review. Under prototype inference, the correct syndrome appeared among the top three candidates for 0.550 of patients, among the top five for 0.652, and among the top ten for 0.781 (Fig. 5a). Prototype inference consistently outperformed softmax inference across candidate list sizes, improving top-5 accuracy by 3.7 percentage points (95% CI, 1.9–5.6) and mean reciprocal rank from 0.462 to 0.490. When each syndrome was weighted equally, macro top-5 recall also increased from 0.539 to 0.593 (Fig. 5b), indicating that the ranking benefit was not limited to frequently represented diagnoses.

**Figure 5:**
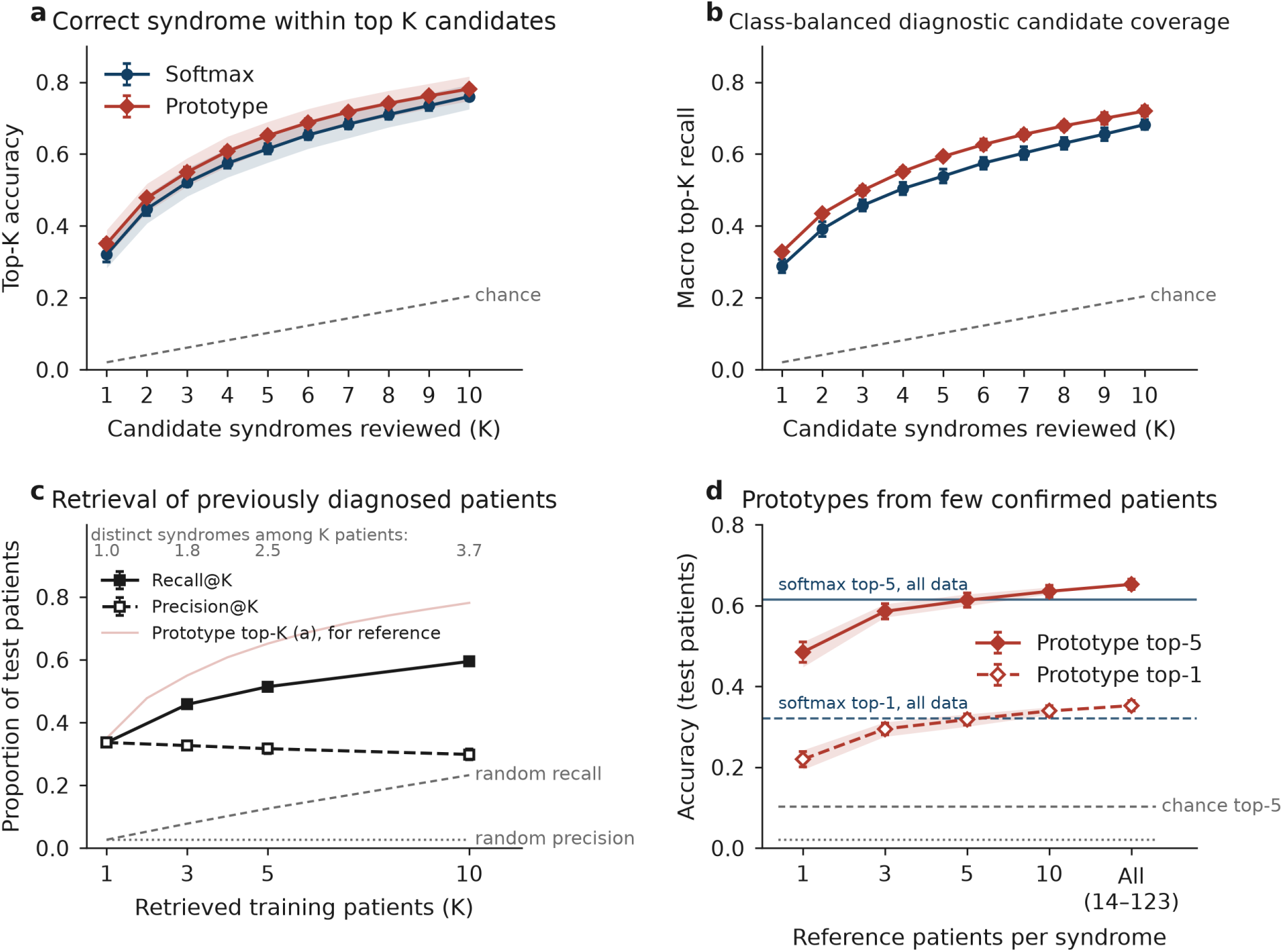
3D facial representations support diagnostic candidate prioritization and patient retrieval. Results for the 49-class syndrome-diagnosis task using five independently trained PointNet++ models without retraining. Each held-out patient was represented by the average of their scan embeddings or predictions. Values are mean *±* SD across trained models, with inference seeds averaged within each model. **(a)** Top-*K* accuracy: proportion of patients whose correct syndrome appeared among the *K* highest-ranked diagnoses under softmax (navy) or prototype (red) inference. Shaded bands indicate 95% patient-level bootstrap confidence intervals; dashed line indicates chance (*K/*49). **(b)** Macro top-*K* recall, calculated by averaging diagnosis-specific top-*K* recall across all 49 syndromes, giving each syndrome equal weight. **(c)** Patient-to-patient retrieval using one averaged embedding per training patient. Recall@*K* is the proportion of queries for which at least one of the *K* nearest patients shared the true diagnosis; Precision@*K* is the mean proportion of retrieved patients sharing that diagnosis. Grey lines indicate random retrieval, and the light red line shows prototype top-*K* accuracy for comparison. Numbers above the panel indicate the average number of distinct syndromes among retrieved patients. **(d)** Top-1 and top-5 accuracy using syndrome prototypes constructed from 1, 3, 5, 10, or all available training patients per diagnosis. The encoder remained fixed. Shaded bands indicate the 2.5th–97.5th percentile across 50 random reference selections; error bars indicate SD across trained models. Horizontal lines show softmax performance using all training data and chance levels.

We next evaluated whether the learned representations could retrieve previously diagnosed patients with similar facial morphology. Using one averaged embedding per training patient, the nearest retrieved patient shared the query patient’s diagnosis for 0.336 of held-out patients, increasing to 0.514 and 0.595 when considering the five and ten nearest patients, respectively (Fig. 5c). Approximately one-third of retrieved patients shared the query diagnosis, substantially exceeding random retrieval. However, the five nearest patients represented only 2.5 distinct syndromes on average, resulting in lower diagnostic coverage than ranking five syndrome prototypes. These results suggest complementary roles for the two approaches: prototype inference prioritizes candidate diagnoses, whereas patient-level retrieval identifies phenotypically similar confirmed cases for comparison.

Finally, we investigated whether useful prototypes could be constructed from limited numbers of reference patients. Keeping the encoder fixed, we constructed syndrome prototypes from 1, 3, 5, or 10 randomly selected training patients per diagnosis. With only one reference patient per syndrome, prototype inference achieved top-1 accuracy of 0.220 and top-5 accuracy of 0.485 (Fig. 5d). Performance improved substantially with the first few reference patients, with three patients per syndrome recovering 90% of the top-5 accuracy obtained using all available training patients. With five reference patients, top-5 accuracy reached 0.613, comparable to the softmax classifier using all training data (0.615). These findings indicate that the learned embedding space supports diagnostic prioritization using relatively small reference libraries, although the encoder had been trained on all 49 syndromes.

The results support a retrieval-oriented role for 3D facial representations in differential diagnosis. Prototype-based ranking can prioritize a short list of candidate syndromes, while patient-to-patient retrieval provides complementary examples of confirmed cases. However, these findings reflect evaluation within a predefined set of diagnoses and do not establish recognition of previously unseen syndromes.

### 2.6 Diagnostic performance varies across syndromes beyond class size

Performance varied substantially across individual diagnoses (Fig. 6). Diagnoses with more test subjects tended to achieve higher per-class *F*_1_ (Spearman *ρ* = +0.35, 95% CI [+0.07, +0.60]), indicating a positive association between class size and diagnostic performance. However, substantial variation remained among diagnoses with similar numbers of test subjects, suggesting that class size alone does not determine how readily a syndrome can be distinguished from 3D facial morphology.

**Figure 6:**
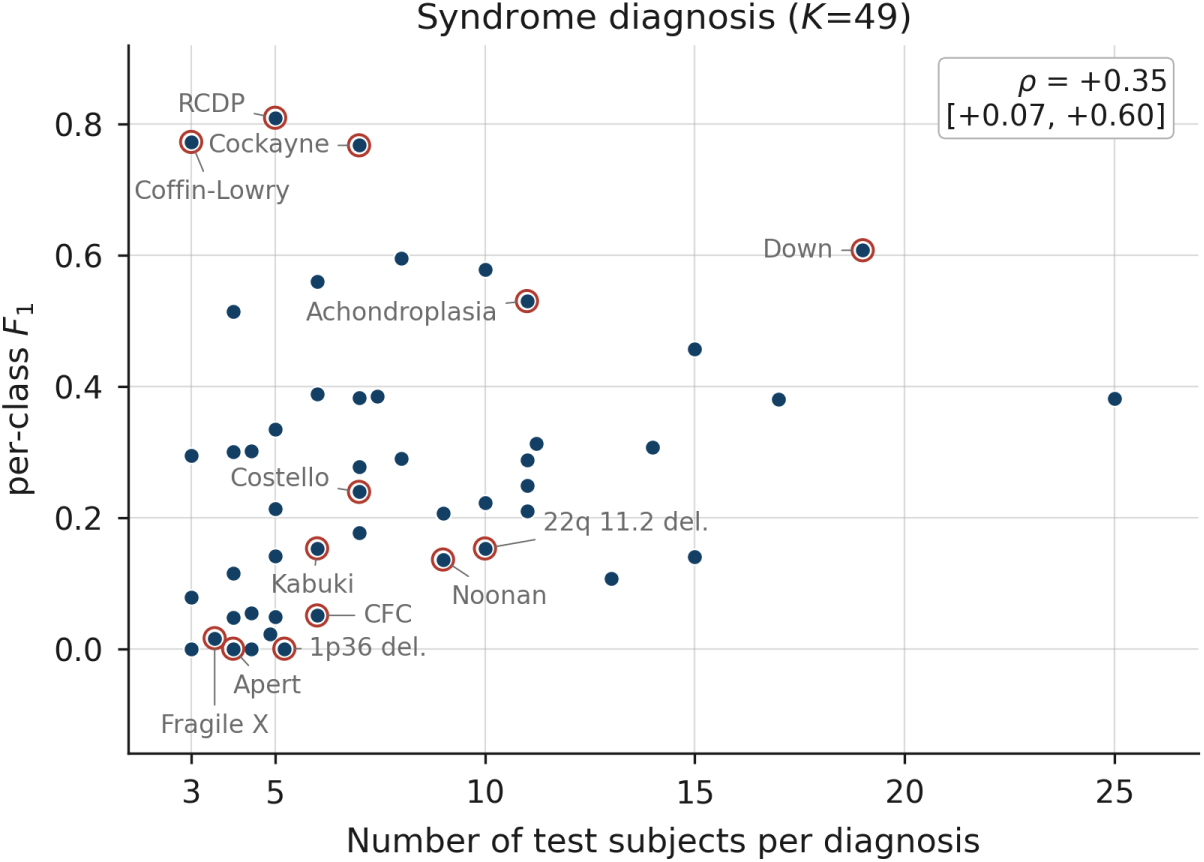
Association between class size and diagnostic performance. Per-class *F*_1_ is plotted against the number of test subjects for the 49-class syndrome-diagnosis task. Values represent subject-level PointNet++ performance averaged across five independently trained models, with inference-seed variation averaged within each training run. Each point represents one diagnosis, and the *x*-axis gives the number of test subjects with that diagnosis. The inset reports the Spearman correlation (*ρ*) between class size and per-class *F*_1_, with 95% bootstrap confidence intervals obtained by resampling classes. Larger classes tended to achieve higher *F*_1_, although substantial performance variation remained among diagnoses with similar class sizes. Selected diagnoses discussed in the text are circled and labelled. RCDP, rhizomelic chondrodysplasia punctata; CFC, cardiofaciocutaneous syndrome.

Sensitivity analysis across alternative class-size thresholds further separated the effect of training-set inclusion from the difficulty introduced by a larger diagnostic label space. Increasing the minimum number of subjects required per diagnosis reduced the number of candidate diagnoses and increased overall top-1 accuracy, whereas macro AUC changed more modestly (Supplementary Fig. S4,b). When evaluation was restricted to the same 12 diagnoses and the same 173 test subjects across all thresholds, accuracy remained nearly constant despite differences in the number of subjects used to define the training task (Supplementary Fig. S4). Thus, much of the increase in top-1 accuracy at stricter thresholds reflected the smaller number of competing diagnoses rather than improved recognition of the diagnoses shared across thresholds.

Marked diagnosis-specific differences nevertheless remained at comparable class sizes (Fig. 6). Among the 24 diagnoses represented by 3–6 test subjects, per-class *F*_1_ ranged from 0.00 to 0.81. More broadly, sparsely represented diagnoses including rhizomelic chondrodysplasia punctata, Coffin– Lowry syndrome, and Cockayne syndrome achieved *F*_1_ values of 0.77–0.81, whereas Apert syndrome, Fragile X syndrome, and 1p36 deletion showed little or no correct classification at similar class sizes. Down syndrome, despite having one of the largest test sets, achieved an intermediate *F*_1_ of approximately 0.61. These results indicate that additional diagnosis-specific factors contribute to recognizability beyond the number of available examples.

Classification errors were also concentrated in particular alternative diagnoses. For each syndrome with at least two misclassified subjects, we identified the most frequently predicted incorrect diagnosis and calculated its proportion of all errors for that syndrome. On average, 0.35 of errors were assigned to the most common alternative diagnosis across 47 syndromes, indicating recurring patterns of diagnostic confusion. We further examined whether errors were preferentially concentrated among phenotypically related syndromes. Among misclassified subjects with Noonan syndrome, Costello syndrome, or cardiofaciocutaneous syndrome (CFC), 17% were assigned to another RASopathy, compared with a RASopathy prevalence of 6% in the test set. This corresponded to approximately 2.9-fold enrichment, and the within-family fraction ranged from 6% to 33% across the five independently trained models (Supplementary Fig. S5). The enrichment was driven primarily by CFC subjects predicted as Costello syndrome. Nevertheless, most RASopathy errors (83%) were assigned to diagnoses outside the group. These findings show that clinically related syndromes can exhibit preferential confusion, although syndrome-family similarity accounts for only a subset of the remaining diagnostic errors.

### 2.7 Anatomical interpretability of learned 3D facial representations

To investigate whether the learned 3D facial representations capture clinically meaningful craniofacial characteristics, we performed an anatomical region-occlusion analysis using the five PointNet++ models trained for the 49-class syndrome-diagnosis task (Methods). Facial surfaces were partitioned into seven regions using automatically detected facial landmarks: forehead, periorbital, nose, cheek, philtrum, lips, and chin/jaw. Regional importance was quantified as the reduction in similarity to the true-diagnosis prototype following region removal, relative to a randomly selected facial patch of matched size.

Representative subjects correctly classified by all five models revealed distinct patterns of regional importance (Fig. 7a). In achondroplasia, the forehead and nasal regions contributed most strongly, consistent with characteristic frontal bossing and nasal–midfacial abnormalities. Down syndrome showed prominent contributions from the cheek and periorbital regions, whereas Pallister–Killian syndrome exhibited broader importance across the cheek, periorbital, nasal, and forehead regions. Cockayne syndrome showed prominent periorbital importance, while rhizomelic chondrodysplasia punctata (CDP) exhibited its strongest contribution in the periorbital region, followed by smaller contributions from the lips, cheeks, and nose. These patterns broadly aligned with clinically recognized craniofacial features of the respective syndromes, suggesting that the learned representations capture anatomically meaningful characteristics.

**Figure 7:**
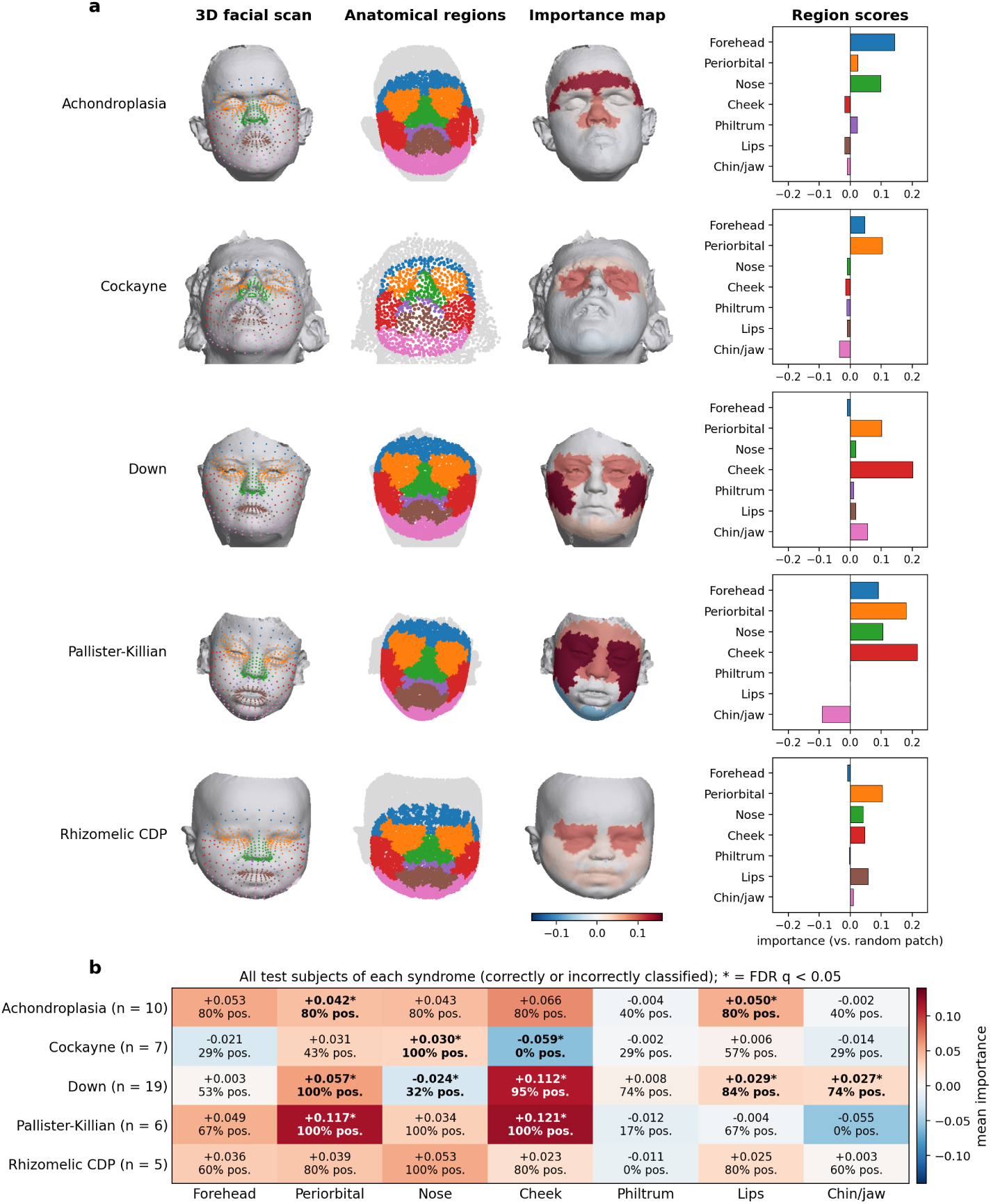
Anatomical interpretability of learned 3D facial representations across genetic syndromes. **(a)** Representative subjects correctly classified by all five PointNet++ models. For each syndrome, the 3D facial scan with automatically detected landmarks, anatomical regions of the 4,096-point model input, regional importance map, and corresponding importance scores are shown. Seven facial regions were defined: forehead, periorbital, nose, cheek, philtrum, lips, and chin/jaw. Importance was measured as the reduction in cosine similarity to the true-diagnosis prototype after region removal, relative to removing a random facial patch of matched size. Positive scores indicate greater importance than the random-patch baseline. Importance maps share a fixed color scale of *±*0.16; gray regions fall outside the anatomical masks. **(b)** Mean regional importance across all test subjects within each syndrome, including correctly and incorrectly classified cases. Cells show the mean importance score and percentage of subjects with positive importance, with sample sizes (*n*) indicated for each syndrome. Asterisks and bold text indicate regions with statistically significant importance (two-sided one-sample *t*-test, Benjamini–Hochberg false discovery rate (FDR) correction *q <* 0.05 across the 35 syndrome–region comparisons shown).

To examine whether these patterns generalized beyond individual examples, we aggregated regional importance across all test subjects within each syndrome, including both correctly and incorrectly classified cases (Fig. 7b). Down syndrome showed strong cheek (+0.112) and periorbital (+0.057) importance, positive in 95% and 100% of subjects, respectively, with both regions remaining significant after FDR correction. Similarly, Pallister–Killian syndrome exhibited significant cheek (+0.121) and periorbital (+0.117) contributions across all six subjects, indicating consistent reliance on these facial regions. Achondroplasia showed a more distributed pattern involving the forehead, nose, cheeks, periorbital region, and lips, each positive in 80% of subjects, although only the periorbital and lip regions remained significant after correction. Nasal importance was positive in all subjects with Cockayne syndrome and rhizomelic CDP, but reached significance only in Cockayne syndrome. Notably, Cockayne syndrome also showed significant negative relative cheek importance and variable periorbital contributions, illustrating how regional reliance can differ between individual subjects and broader syndrome group.

Overall, the learned 3D facial representations showed anatomically meaningful regional importance, with recurring patterns in some syndromes and greater variability in others. However, these regions contribute to true-diagnosis similarity and are not necessarily specific to individual syndromes.

Nearest-neighbor examples further illustrated syndrome-associated facial similarity among correctly classified subjects (Supplementary Fig. S6). Nevertheless, misclassified cases (Supplementary Fig. S7) revealed confusion between clinically related conditions, including cardiofaciocutaneous and Costello syndromes and Marfan and Loeys–Dietz syndromes, highlighting the challenges of distinguishing syndromes with similar facial characteristics.

### 2.8 Comparison with published 3D craniofacial approaches

We contextualized our findings with previously published 3D craniofacial analysis methods (Table 4). Because these studies differ in cohort composition, diagnostic categories, preprocessing, train–test partitioning, and evaluation metrics, their reported performance values are not directly comparable.

**Table 4:** Comparison with previously published 3D craniofacial analysis methods. Reported performance values are shown together with the use of registration, number of classes, and evaluation metric for each study. Because cohorts, diagnostic categories, data splits, and performance metrics differ across studies, these results provide contextual rather than controlled comparisons. Values are reproduced from the corresponding publications.

| Study | Registration | Cls | Metric | Value |
|---|---|---|---|---|
| *Registered pipelines* |  |  |  |  |
| Hallgrímsson et al. [13] | dense correspondence | 64 | top-1, syndromic subj. | 48.8% |
|  |  |  | top-10, same cohort | 87.2% |
| Bannister et al. [17] | statistical shape model | 48 | overall top-1 (incl. unaff.) | 71% |
|  |  |  | mean syndrome sensitivity | 43% |
| Bannister et al. [35] | statistical shape model | 2 | ROC-AUC, syndrome vs. normal | 0.863 |
| *Unregistered pipelines* |  |  |  |  |
| Bannister et al. [11] | none (3D PCA + MLP) | 43 | mean top-1 sensitivity | 40.7% |
| *This work* (subject-level evaluation, no registration) |  |  |  |  |
| PointNet++, softmax | none | 49 | top-1, syndromic subj. | 0.321 |
| PointNet++, prototype | none | 49 | top-1, syndromic subj. | 0.350 |
| PointNet++, prototype | none | 49 | top-5, syndromic subj. | 0.652 |
| PointNet++, softmax | none | 2 | ROC-AUC, affected vs. unaff. | 0.956 |

For broad affected-versus-unaffected screening, our PointNet++ model achieved an AUC of 0.956, whereas a registered statistical shape-model approach reported an AUC of 0.863 for syndrome-versus-normal discrimination [35]. Our cohort sensitivity analyses also showed that combined screening performance was substantially influenced by cohort composition and demographic differences (Section 2.3), highlighting the importance of accounting for population characteristics when interpreting screening results.

For fine-grained syndrome diagnosis, our patient-level PointNet++ model achieved top-1 accuracy of 32.1% with softmax inference and 35.0% with prototype inference, with prototype top-5 accuracy reaching 0.652. An unregistered 3D PCA-plus-MLP approach reported mean top-1 sensitivity of 40.7% across 43 classes [11]. Registered correspondence-based approaches have also reported higher performance on some syndrome-recognition tasks. For example, Hallgŕımsson et al. [13] reported top-1 accuracy of 48.8% across 64 syndromes, while Bannister et al. [17] reported mean syndrome sensitivity of 43% across 48 syndrome classes. These differences cannot be attributed to registration or model architecture without controlled comparisons using the same cohorts and evaluation procedures.

Overall, our study extends existing 3D craniofacial benchmarks by systematically evaluating multiple facial representations across screening and fine-grained diagnostic tasks. The additional analyses of patient-level aggregation, prototype-based candidate ranking, and retrieval of confirmed cases further highlight the potential of 3D facial representations and extend conventional classification benchmarks toward clinically relevant diagnostic prioritization.

## 3 Discussion

### 3.1 Registration-free representations capture clinically relevant morphology

Our results show that craniofacial morphology associated with congenital and genetic conditions can be learned directly from 3D facial surface geometry without landmark registration, dense correspondence, or fitting individual faces to a normative template. Across multiple architectures and clinical tasks, these representations captured disease-associated facial differences, although performance varied substantially with diagnostic specificity and cohort composition.

This registration-free framework offers several practical advantages. It simplifies preprocessing, avoids dependence on manually or automatically detected landmarks, and eliminates the need to deform each face toward a common template. These properties may be especially valuable for craniofacial disorders, where substantial anatomical abnormalities can make landmarking and template fitting more difficult and may reduce measured deviations from normative facial shape. By operating directly on sampled surface geometry, the models retain native facial morphology while providing a common representation that can be applied across heterogeneous craniofacial phenotypes.

More broadly, these findings demonstrate that dense anatomical correspondence is not required to recover disease-associated information from 3D facial scans. This provides a flexible approach to craniofacial phenotyping, particularly in heterogeneous cohorts where consistent anatomical registration may be challenging. However, establishing the relative advantages of registration-free and correspondence-based approaches will require direct comparisons using the same patient populations and evaluation tasks.

### 3.2 From screening to differential diagnosis

The dominant pattern across experiments was a decline in performance as greater diagnostic specificity was required. Broad craniofacial screening achieved high performance in the combined benchmark, whereas cleft subtyping and syndrome-level diagnosis were substantially more difficult. This pattern was consistent across model architectures, suggesting that distinguishing increasingly subtle and overlapping craniofacial phenotypes presents a substantial challenge beyond the choice of a particular representation learning model.

Importantly, our cohort sensitivity analyses showed that high combined-screening performance was strongly influenced by cohort composition and demographic differences. Source-cohort membership alone was highly predictive of affected status, and discrimination decreased in within-cohort and age- and sex-matched comparisons. Although facial representations retained discriminatory information after accounting for these factors, the limited availability of unaffected pediatric subjects, particularly in the syndrome cohort, restricted the assessment of screening performance in clinically relevant populations. These findings highlight the importance of distinguishing disease-associated facial morphology from demographic and cohort-related variation when evaluating AI-based craniofacial screening.

Fine-grained syndrome diagnosis presented a different challenge. Class-wise discrimination remained comparatively strong despite limited top-1 accuracy, and the correct diagnosis was frequently retained among the highest-ranked candidates. Performance also varied substantially among syndromes with similar class sizes, while related RASopathies showed preferential diagnostic confusion. These observations suggest that syndrome recognizability depends not only on the amount of available training data but also on the distinctiveness, heterogeneity, and overlap of facial phenotypes.

Repeated scans provided additional information for fine-grained diagnosis. Our controlled analysis showed that improvements from patient-level evaluation reflected both differences in patient weighting and the additional benefit of combining scans. The largest gains generally occurred when combining the first few scans, particularly under prototype inference, although individual patients did not always benefit. This suggests that repeated acquisitions can provide complementary information about facial morphology when diagnostic differences are subtle. It also emphasizes that the patient, rather than the individual scan, is the clinically relevant unit of inference. Together with the importance of subject-disjoint data partitioning, these findings highlight how evaluation design and patient-level representation affect the interpretation of diagnostic performance.

### 3.3 Prototype inference supports a retrieval-oriented clinical use case

Prototype inference provided a clinically relevant alternative to conventional softmax classification. Although it produced modest improvements in top-1 accuracy and macro-*F*_1_ without substantially changing class-wise discrimination, its principal value was in prioritizing candidate diagnoses. The correct syndrome appeared among the five highest-ranked candidates for nearly two-thirds of held-out patients, and prototype inference consistently improved ranking compared with softmax inference, including when diagnoses were weighted equally. This distinction is relevant to clinical genetics, where facial morphology is typically used to narrow a differential diagnosis and guide further clinical or genetic evaluation rather than establish a definitive diagnosis from facial appearance alone.

Syndrome-level prototype ranking and patient-to-patient retrieval offered complementary information. Comparing a patient’s facial representation with syndrome prototypes provided a more diverse list of candidate diagnoses, whereas retrieving previously diagnosed patients identified phenotypically similar confirmed cases that could support expert review. Moreover, prototype ranking remained informative when only a few reference patients per syndrome were available, suggesting potential utility in settings where confirmed cases are limited. However, the encoder was trained on all 49 syndromes, so this finding does not establish the ability to recognize previously unseen conditions.

These results support a retrieval-oriented role for 3D facial representations as a potential complement to clinical and genomic evaluation. Their clinical utility will require further validation in representative patient populations, particularly when the correct diagnosis is not included in the reference library.

### 3.4 Anatomical interpretability of learned facial representations

Anatomical interpretability analysis provided further evidence that the learned 3D facial representations capture clinically meaningful craniofacial morphology. Region occlusion revealed distinct patterns of facial importance across genetic syndromes, several of which aligned with clinically recognized features. For example, achondroplasia showed contributions from the forehead and nasal regions, consistent with frontal bossing and midface abnormalities, whereas Down syndrome and Pallister–Killian syndrome showed prominent cheek and periorbital importance. These observations suggest that the models capture anatomically meaningful facial characteristics rather than relying solely on global facial shape.

Importantly, syndrome-level analysis revealed that some regional importance patterns were consistent across individuals, while others showed substantial variability. The cheek and periorbital regions contributed consistently in Down syndrome and Pallister–Killian syndrome, whereas Cockayne syndrome exhibited more variable periorbital importance despite its prominence in the representative example. Such variability may reflect differences in facial expression, age, phenotype severity, or individual morphology, although these factors were not directly evaluated. Moreover, regions important for representing a particular syndrome are not necessarily specific to that diagnosis, as different conditions may share similar facial characteristics.

This distinction was further illustrated by misclassified subjects. Confusion between clinically related conditions, including cardiofaciocutaneous and Costello syndromes and Marfan and Loeys– Dietz syndromes, highlights the difficulty of distinguishing similar craniofacial phenotypes. Some misclassified subjects also retained positive importance in clinically relevant regions or retrieved correctly labeled training subjects, indicating that meaningful anatomical information does not always translate into correct diagnostic assignment. Region occlusion therefore provides insight into the anatomical basis of learned representations while also revealing limitations in their ability to distinguish closely resembling conditions.

### 3.5 Limitations and future directions

Several limitations should be considered. Many diagnostic classes remained small despite the minimum inclusion threshold, limiting the precision of class-specific estimates and excluding some of the rarest conditions. The unaffected cohorts were heterogeneous, ancestry information was unavailable for several datasets, and family identifiers were not available to explicitly control for relatedness. Cohort composition and demographic differences also influenced combined screening performance, while the scarcity of unaffected pediatric subjects limited evaluation of syndrome screening in children. In addition, most affected cleft scans were acquired after the typical age of primary repair, and surgical status was not consistently available at the scan level. Consequently, the findings may predominantly reflect postoperative facial morphology and may not generalize directly to unrepaired clefts or preoperative screening.

The models relied exclusively on 3D facial surface geometry, without incorporating complementary information such as age, sex, facial texture, clinical findings, or genomic data. Such information may be particularly valuable for distinguishing syndromes with similar or variable facial characteristics. Although region-occlusion analysis identified anatomically meaningful patterns, it measured regional contributions to true-diagnosis similarity rather than directly explaining individual classification decisions. Furthermore, the relatively small number of subjects in some syndromes limits the assessment of consistency across individuals, and the results do not establish whether the highlighted regions are uniquely discriminative for each diagnosis.

Our diagnostic prioritization analyses were also restricted to the 49 syndromes represented during encoder training. Thus, performance with limited reference patients does not demonstrate generalization to previously unseen syndromes, and diagnostic ranking was not evaluated when the correct diagnosis was absent from the candidate library. Patient-level aggregation was assessed using repeated acquisitions without information on acquisition timing or clinical changes between scans. Whether similar benefits would be observed with prospectively collected or longitudinal scans remains uncertain.

Future work should prioritize larger and more demographically diverse cohorts, particularly age-matched pediatric controls, and approaches tailored to rare and imbalanced diagnoses. Evaluating diagnostic prioritization in broader reference libraries, including previously unseen syndromes and patients without a matching diagnosis, will be important for establishing clinical applicability. Integration of facial morphology with clinical and genomic information, together with prospective evaluation of patient-level inference and expert-assisted diagnostic workflows, may further clarify the clinical utility of these approaches.

Overall, our findings demonstrate that 3D facial representations can capture disease-associated morphology across a spectrum of craniofacial conditions, although screening performance requires careful consideration of cohort and demographic differences. Fine-grained syndrome diagnosis remains challenging, but patient-level aggregation, prototype-based candidate ranking, and retrieval of confirmed cases provide complementary approaches to diagnostic prioritization. These capabilities support further development of 3D facial analysis as an aid to clinical and genetic evaluation.

## 4 Methods

### 4.1 Cohorts and datasets

All 3D facial data were obtained from the FaceBase consortium repository. Six datasets were included, comprising two orofacial cleft cohorts, one genetic-syndrome cohort, and three unaffected comparison cohorts (Table 1, Fig. 1b).

**Orofacial cleft cohorts.**

FB-5A and FB-56 are family-based studies of non-syndromic orofacial clefting recruited across multiple international sites. Both datasets provide 3D facial surfaces as OBJ meshes, together with subject identifiers and phenotype information. Facial surfaces were linked to participant-level cleft labels using dataset-specific image-mapping and participant tables. Scans without a recorded Cleft Type were excluded. The cohorts included both pre- and postoperative scans, although most affected scans were acquired after the typical age of primary cleft repair: 88% (1,090/1,238) were obtained at age *≥*2 years, whereas approximately 4–6% were acquired before 6 months. Available surgical history also indicated that repaired cases outnumbered unrepaired cases. After preprocessing, FB-5A contributed 3,099 scans and FB-56 contributed 3,839 scans.

### Genetic-syndrome cohort

FB-TJ0 (FaceBase accession FB00000861) is a prospective 3D facial phenotype collection of individuals with genetic syndromes and their family members. Facial surfaces were acquired using 3dMD photogrammetry systems and distributed as PLY files, with metadata including subject identifiers, acquisition dates, demographic characteristics, clinical findings, diagnoses, syndrome names, OMIM numbers, and syndrome categories. FB-TJ0 was the only included cohort with diagnosis-level annotations for genetic syndromes. A total of 13,307 scans were successfully processed.

### Unaffected comparison cohorts

FB-TK0, FB-TX4, and FB-VWP contained unaffected individuals and contributed 686, 3,605, and 2,454 scans, respectively. These cohorts lacked diagnostic phenotype labels and were therefore used only for the combined affected-versus-unaffected screening task. Each subject contributed a single scan.

Repeated imaging occurred primarily in FB-TJ0, where subjects contributed a median of 2.6 scans and up to 34 scans per individual. The cleft and unaffected comparison cohorts contained approximately one scan per subject. This repeated-measures structure was accounted for during data partitioning and patient-level evaluation.

### 4.2 Clinical tasks and label definitions

We defined four prediction tasks spanning broad screening to fine-grained diagnosis: combined affected-versus-unaffected screening, binary cleft screening, four-class cleft subtype classification, and 49-class genetic syndrome diagnosis (Fig. 1c).

### Cleft tasks

Subjects in FB-5A and FB-56 were labeled as Unaffected, Cleft Lip, Cleft Palate, or Cleft Lip and Palate. Records labeled “Unknown Cleft” were excluded because they indicated clefting without a specific subtype. These four labels defined the cleft-subtyping task. For binary cleft screening, the three cleft categories were combined into a single affected class and compared with unaffected subjects.

### Syndrome-diagnosis task

Labels were obtained from the FB-TJ0 *Syndrome* field, which specifies the diagnosed condition for each subject. Values differing only in capitalization were merged under their most frequent spelling. Entries that did not identify a single confirmed disorder, including Relative, Unknown, and “Facial Dysmorphia, possibly genetic,” were excluded. When the independently recorded *Clinical Diagnosis* field also named a disorder, it was compared with the Syndrome field; subjects with discordant diagnoses were excluded (21 subjects and 39 scans).

Diagnoses were retained if they contained at least 50 scans and 20 subjects, yielding 49 diagnoses comprising 7,472 scans from 2,719 subjects (20–173 subjects per diagnosis). The inclusion threshold was fixed before model training. Sensitivity to this threshold was evaluated by rebuilding the task at minimum class sizes of 10, 15, 30, 40, 50, and 75 subjects, corresponding to 55 to 12 retained diagnoses. A common subject-level split was assigned within each diagnosis so that test sets were nested across thresholds, and all four models were retrained with five seeds at each threshold.

### Combined screening task

All cleft subtypes and labeled FB-TJ0 syndrome cases were assigned to the affected class. Unaffected subjects from the cleft cohorts, the FB-TJ0 Control group, and the three unaffected comparison cohorts were assigned to the unaffected class. The resulting dataset contained 24,973 scans.

### 4.3 3D facial surface preprocessing

All datasets were processed using the same geometry-only pipeline. OBJ and PLY surfaces were loaded without texture or material information, retaining only vertex coordinates and mesh geometry. Coincident vertices were merged to avoid artificial separation of surfaces at texture seams. Connected components were filtered by retaining the component with the largest surface area and iteratively adding components located within 4% of the mesh bounding-box diagonal of the retained surface. This procedure preserved adjacent components produced by multi-camera facial capture while removing detached fragments and distant background geometry.

Triangle meshes were initially sampled to approximately three times the target point count using surface-area-based sampling. Poisson-disc-style sampling was used when available, with area-weighted sampling as a fallback. The resulting point set was reduced to *N* = 4,096 points using farthest-point sampling. PLY files containing point sets rather than mesh faces were downsampled directly using farthest-point sampling.

Each point was centered at its centroid and isotropically scaled so that the maximum Euclidean distance from the centroid was one. No facial landmarking, Procrustes alignment, dense correspondence, template-based cropping, or rigid or non-rigid template registration was applied. The pipeline therefore normalized translation and global scale while retaining native facial surface geometry.

### 4.4 Model architectures

We evaluated three point-cloud neural networks and one handcrafted geometric baseline: Point-Net [25], PointNet++ [26], DGCNN [27], and GeomMLP. All approaches used the same preprocessed 4,096-point facial representation, either directly or through extracted geometric descriptors. The architectures are summarized in Fig. 8.

**Figure 8:**
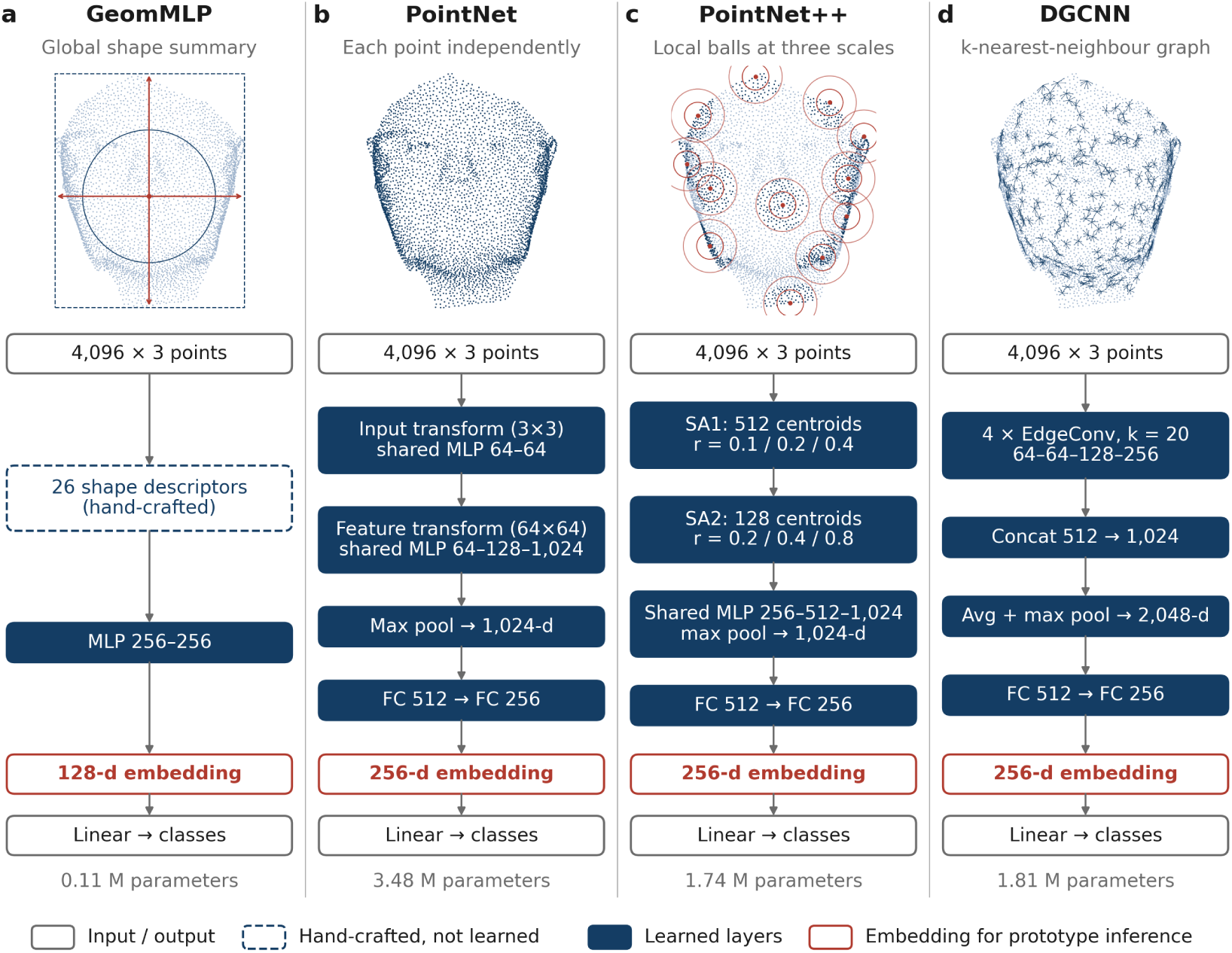
Model architectures for 3D facial representation learning. Each approach uses the same preprocessed 4,096-point facial point cloud. Upper panels illustrate the geometric information used, and lower panels summarize the architectures. **(a)** GeomMLP extracts 26 handcrafted global shape descriptors, including principal-axis, bounding-box, and radial-distance features, followed by a multilayer perceptron. **(b)** PointNet applies shared point-wise transformations and multilayer perceptrons followed by global max pooling. **(c)** PointNet++ uses multi-scale local neighborhoods around centroids selected by farthest-point sampling in two set-abstraction stages. **(d)** DGCNN constructs dynamic *k*-nearest-neighbor graphs (*k* = 20) and extracts local features through EdgeConv layers. The three point-cloud models use a common classification head with 512- and 256-unit fully connected layers. Red boxes indicate the penultimate representations used for prototype inference (256 dimensions for point-cloud models and 128 dimensions for GeomMLP). Navy boxes denote learned layers, and the dashed box denotes handcrafted features. Parameter counts are shown for each model.

### PointNet

PointNet applies a shared multilayer perceptron independently to each point and aggregates point-wise features through global max pooling. The point-wise network used channel dimensions (64)–(64)–(64)–(128)–(1024). Input and feature transformation networks predicted (3*×*3) and (64*×*64) transformations, respectively, with orthogonality regularization applied to the feature transform. The resulting 1,024-dimensional global descriptor was passed to the classification head. The model contained 3.48 million parameters.

#### PointNet++

We used the multi-scale grouping variant of PointNet++. Two set-abstraction stages reduced the point cloud to 512 and 128 centroids using farthest-point sampling. At the first stage, local neighborhoods were defined using radii of 0.1, 0.2, and 0.4 with maximum neighborhood sizes of 16, 32, and 128 points. At the second stage, radii of 0.2, 0.4, and 0.8 with maximum neighborhood sizes of 32, 64, and 128 points were used. Features from the three spatial scales were concatenated, followed by global feature aggregation to obtain a 1,024-dimensional descriptor. The model contained 1.74 million parameters.

#### DGCNN

DGCNN used four EdgeConv blocks with 64, 64, 128, and 256 output channels. At each layer, a *k* = 20 nearest-neighbor graph was constructed in the current feature space. Features from all EdgeConv blocks were concatenated and projected to 1,024 channels, followed by concatenated global average and max pooling to obtain a 2,048-dimensional descriptor. The model contained 1.81 million parameters.

### GeomMLP

GeomMLP provided a non-point-cloud baseline using 26 global geometric descriptors derived from each normalized point cloud. These included covariance eigenvalues, summary statistics of coordinates projected onto principal axes, bounding-box dimensions and center, radial-distance statistics, the proportion of points within half the unit radius, and centroid offsets. The descriptors were passed to a three-layer multilayer perceptron with 256, 256, and 128 hidden units. The model contained 0.11 million parameters.

For the three point-cloud encoders, the global representation was followed by a common two-layer classification head with 512 and 256 hidden units, batch normalization, dropout, and a final linear layer over task classes. The 256-dimensional penultimate activation was used as the embedding for prototype-based inference. GeomMLP produced a 128-dimensional penultimate embedding.

### 4.5 Data partitioning and model training

Data were partitioned at the subject level to prevent information leakage from repeated scans. All scans from an individual were assigned to the same partition. Subject identifiers were constructed by combining the dataset identifier with the dataset-specific participant identifier because some identifiers were reused across FaceBase cohorts. Subjects whose repeated scans carried inconsistent labels were excluded before partitioning.

Within each class, subjects were randomly shuffled using a fixed seed and divided into training, validation, and test sets using a 70%/15%/15% split, with at least one subject assigned to the validation and test sets when possible. For the syndrome-diagnosis task, this yielded 1,949, 385, and 385 subjects (5,308, 1,108, and 1,056 scans), with at least 3 test subjects per diagnosis.

All models were optimized with AdamW using an initial learning rate of 10*^−^*^3^ and weight decay of 10*^−^*^4^. A cosine-annealing schedule reduced the learning rate to 10*^−^*^5^ over 200 epochs. Batch size was 16 for PointNet++ and 32–64 for the remaining models. Training used class-weighted cross-entropy, with weights inversely proportional to class frequency in the training set.

Models were trained for the full 200 epochs. Early stopping was not used because preliminary experiments showed extended optimization plateaus for several tasks, causing patience-based stopping to terminate some runs before learning had stabilized. For each run, the checkpoint with the highest validation macro-*F*_1_ was selected for test evaluation.

Unless otherwise specified, each model–task configuration was trained independently with five random seeds while keeping the data partition fixed.

### 4.6 Inference strategies and patient-level aggregation

We evaluated two inference strategies using the learned facial representations: the trained softmax classifier and prototype inference (Fig. S1e). Both were applied without modifying the trained model weights.

For softmax inference, each scan was assigned class probabilities by the trained classifier, and the highest-probability class was selected as the top-1 prediction. For prototype inference, the final classification layer was removed at inference, and the penultimate embedding was used instead. A prototype for each diagnosis was constructed by averaging the embeddings of all training scans belonging to that class. Test embeddings were compared with the prototypes using cosine similarity, and the most similar diagnosis was selected.

Predictions were evaluated at both scan and patient levels. At scan level, each acquisition was evaluated independently. At patient level, repeated scans from the same individual were combined before classification. For softmax inference, class probabilities were averaged across scans; for prototype inference, scan embeddings were averaged before comparison with class prototypes.

### 4.7 Performance metrics and statistical analysis

Model performance was evaluated using top-1 accuracy, top-5 accuracy, macro-*F*_1_, and macro one-versus-rest AUC. Top-1 accuracy was the proportion of samples for which the correct class received the highest score, whereas top-5 accuracy was the proportion for which the correct class appeared among the five highest-scoring classes. Chance accuracy was 1*/K* for top-1 and 5*/K* for top-5, where *K* is the number of classes. Macro-*F*_1_ was calculated independently for each class and averaged across classes. For multiclass AUC, each class was treated in turn as the positive class against all remaining classes, and the resulting one-versus-rest AUCs were macro-averaged. Chance AUC was 0.5.

Unless otherwise specified, results are reported as mean *±* standard deviation across five independently trained models using the same data partition. PointNet++ additionally involves stochastic farthest-point sampling during inference. Each trained PointNet++ model was therefore evaluated with five fixed inference seeds, whose results were averaged within each training run before variability across runs was summarized. The other models were deterministic at inference under the implemented pipeline.

Bootstrap resampling was used for selected statistical comparisons. Paired patient-level comparisons used the same resampled patients for both configurations, preserving the paired evaluation design. Confidence intervals for the association between class size and per-class *F*_1_ were obtained by bootstrapping diagnostic classes. Class-specific performance was calculated within each model before averaging across training runs. Associations between class size and per-class *F*_1_ were evaluated using Spearman correlation. Additional statistical procedures specific to individual analyses are described below.

### 4.8 Cohort-confounding and demographic sensitivity analyses

To assess the influence of cohort composition and demographic differences on screening performance, we conducted sensitivity analyses using patient-level predictions from the five previously trained PointNet++ models for the combined affected-versus-unaffected task, without retraining or recalibration. Patient-level probabilities were obtained by averaging softmax predictions across repeated scans. Age at scan and sex were obtained from cohort metadata where available (FB-5A, FB-56, FB-TJ0, and FB-VWP); analyses requiring these variables were restricted to subjects with complete information. Age and sex were unavailable for FB-TK0 and FB-TX4.

### Cohort-identity baseline

Each test subject was assigned a score equal to the prevalence of affected subjects among training and validation subjects from the same source cohort. Discrimination was evaluated using patient-level AUC on the original held-out test set. This baseline did not use facial or demographic information.

### Within-cohort evaluation

Existing PointNet++ predictions were evaluated separately within FB-TJ0, FB-5A, and FB-56, as well as within the two cleft cohorts combined. For each subset, AUC was calculated using the corresponding affected and unaffected test subjects. We also calculated the false-positive rate among unaffected test subjects in each cohort, defined as the proportion assigned to the affected class using the original probability threshold of 0.5.

### Demographic comparison models

To quantify the predictive contribution of age, we fitted an age-only logistic regression model using age and log(1 + age) as predictors. Models were trained on the corresponding non-test subjects and evaluated on held-out test subjects within FB-TJ0 and the combined cleft cohorts. To investigate whether facial predictions contributed information beyond age and sex, we compared two additional logistic regression models: one using age, log(1 + age), and sex, and another additionally incorporating the logit-transformed PointNet++ affected probability. These models were evaluated using stratified five-fold cross-validation repeated 20 times within the held-out test subjects. AUC was calculated from out-of-fold predictions and averaged across repetitions. This secondary analysis evaluated the incremental information provided by the existing facial scores without retraining the PointNet++ models.

### Age-restricted and matched evaluation

Screening performance was separately evaluated among subjects younger than 18 years within FB-TJ0 and the cleft cohorts. We also performed 1:1 age- and sex-matched comparisons within source cohorts. Affected test subjects were processed in a fixed random order and matched without replacement to the closest-age unaffected subject of the same sex, requiring an age difference of no more than two years. Affected subjects without an eligible match were excluded. The same matched subjects were used across the five PointNet++ models. The resulting samples included 147 matched pairs in the cleft cohorts (median affected age, 8.3 years) and 107 pairs in FB-TJ0 (median affected age, 31.0 years).

### Uncertainty estimation

For selected analyses, we additionally constructed an ensemble by averaging patient-level affected probabilities across all 25 model–inference-seed combinations. Ensemble AUC was evaluated using 2,000 patient-level bootstrap resamples, with the 2.5th and 97.5th percentiles defining the 95% confidence interval. The cohort-identity and age-only baselines were reported as single AUC values. Detailed sample sizes and uncertainty estimates are provided in Supplementary Table S1.

### 4.9 Controlled evaluation of repeated-scan aggregation

To distinguish the benefit of combining scans from differences in patient weighting, we evaluated the existing PointNet++ models on fixed groups of patients with repeated scans, without retraining. Within each group of patients with at least *k* scans, predictions were evaluated using *m* = 1*, . . ., k* scans per patient, with each patient contributing equally to performance estimates. For each patient and scan count, results were averaged over all possible subsets of scans or 300 randomly sampled subsets when more were available. We evaluated top-1 accuracy, top-5 accuracy, and the rank of the true syndrome among 49 candidates, where rank 1 indicates the highest-scoring diagnosis.

The contribution of patient weighting was estimated by comparing scan-level accuracy with patient-level accuracy using one scan per patient, averaged across available scans. The additional contribution of scan aggregation was calculated by comparing single-scan patient-level accuracy with performance using all available scans. To examine the effect of adding a second scan, we evaluated all ordered pairs of scans from each patient, comparing the first scan alone with the two scans combined. Changes in top-1 correctness and true-syndrome rank were averaged within each patient before summarizing results across patients.

Analyses were conducted in fixed patient groups with at least two, three, four, or five scans, ensuring that comparisons across scan counts used the same patients. For paired comparisons, 95% confidence intervals were obtained by bootstrapping patient-level differences (2,000 resamples), and two-sided Wilcoxon signed-rank tests were applied to nonzero paired differences. Additional sensitivity analyses of per-patient rank changes are reported in the Supplementary Results. Repeated scans were treated as separate acquisitions without distinguishing imaging sessions or clinical visits.

### 4.10 Diagnostic prioritization and retrieval analyses

We evaluated three complementary diagnostic prioritization analyses using the existing PointNet++ models without retraining. Each of the 385 held-out patients was represented by aggregated scan predictions or embeddings.

### Candidate syndrome ranking

For each patient, the 49 syndromes were ranked by softmax probabilities or cosine similarity to syndrome prototypes. Top-*K* accuracy (*K* = 1*, . . .,* 10) was the proportion of patients whose true diagnosis appeared among the *K* highest-ranked syndromes. Macro top-*K* recall was calculated by averaging syndrome-specific top-*K* recall across the 49 diagnoses, giving each syndrome equal weight. Mean reciprocal rank (MRR) was calculated as the mean reciprocal position of the true diagnosis. Chance top-*K* accuracy was *K/*49.

### Patient-to-patient retrieval

Each of the 1,949 training patients was represented by the mean embedding of their scans, and held-out patients were compared with this reference library using cosine similarity. Recall@*K* was defined as the proportion of queries for which at least one of the *K* nearest training patients shared the true diagnosis, whereas Precision@*K* was the mean proportion of retrieved patients sharing that diagnosis. Random-retrieval baselines were calculated analytically from diagnosis frequencies in the training library, assuming sampling without replacement. Test patients were excluded from the reference library.

### Limited-reference prototype evaluation

To evaluate the influence of reference-library size, syndrome prototypes were constructed from *n ∈ {*1, 3, 5, 10*}* randomly selected training patients per diagnosis, using the mean of their patient-level embeddings. The reference condition used all training patients per syndrome (14–123). Sampling was repeated 50 times, using identical reference selections across models and inference seeds. The encoder remained fixed throughout; this experiment varied only the number of reference patients used to construct each prototype, not the data used to train the representations.

### Statistical comparisons

For limited-reference prototypes, performance was averaged across reference draws, with draw-to-draw variability summarized by the 2.5th–97.5th percentile range. Paired comparisons between inference strategies, patient retrieval and prototype ranking, and limited-reference and full-reference prototypes used patient-level differences averaged across models, inference seeds, and reference draws where applicable. Confidence intervals were obtained from 2,000 patient-level bootstrap resamples, and two-sided Wilcoxon signed-rank tests were applied to nonzero paired differences.

### 4.11 Anatomical interpretability analysis

We performed anatomical region-occlusion analysis using the five independently trained PointNet++ models for the 49-class syndrome-diagnosis task. For each 3D facial scan, 468 landmarks were automatically detected using MediaPipe FaceMesh [36] from a rendered frontal view and mapped onto the normalized point cloud. Seven anatomical regions were defined using these landmarks: forehead, periorbital, nose, cheek, philtrum, lips, and chin/jaw. Each point in the 4,096-point input cloud was assigned to the region of its nearest facial landmark. Landmarks were used only for post-hoc interpretability analysis and were not involved in model training.

To quantify regional importance, all points belonging to an anatomical region were removed, and the point cloud was restored to 4,096 points by randomly duplicating the remaining points. We measured the resulting decrease in cosine similarity between the scan embedding and its true-diagnosis prototype, constructed from training data. As a control, we removed randomly placed facial patches containing the same number of points, repeating the procedure for three patches per region. Regional importance was defined as the decrease in similarity following anatomical region removal minus the mean decrease following random-patch removal. This size-matched baseline helps distinguish anatomical contributions from effects associated with point deletion. Importance scores were averaged across two inference seeds and five models, followed by averaging across scans from the same patient. Positive scores indicate greater importance than the random-patch baseline. Because importance was measured relative to the true diagnosis, it does not directly explain incorrect predictions.

The syndrome-level analysis focused on five well-recognized syndromes: achondroplasia, Cockayne syndrome, Down syndrome, Pallister–Killian syndrome, and rhizomelic chondrodysplasia punctata. Regional importance was summarized across all test patients within each syndrome, regardless of classification correctness. We reported mean importance and the proportion of patients with positive importance. Statistical significance was assessed using two-sided one-sample *t*-tests across patients, with Benjamini–Hochberg false discovery rate correction across the 35 syndrome–region comparisons (*q <* 0.05).

## Author Contributions

T.C. conceived and designed the study, developed and implemented the computational methods, performed the experiments and statistical analyses, interpreted the results, and drafted the manuscript. M.U.A. contributed to study design, data analysis, interpretation of results, and manuscript revision. J.C. and E.C.L. provided clinical expertise in craniofacial disorders and genetic syndromes, contributed to interpretation of the findings, and critically reviewed the manuscript. K.W. supervised the study, contributed to study conception and design, provided scientific guidance, and critically revised the manuscript. All authors read and approved the final manuscript.

## Data Availability

The 3D facial imaging datasets analyzed in this study were obtained from the FaceBase repository (https://www.facebase.org) and are available subject to applicable data access requirements. The datasets include FB-5A [29], *Pittsburgh Oral Facial Cleft Studies: Extending the Phenotype of Nonsyndromic Orofacial Clefting (OFC1)* (FB00001369); FB-56 [30], *Oral-Facial Cleft Families: Phenotype and Genetics (OFC2)* (FB00001368); FB-TJ0 [31], *Developing 3D Craniofacial Morphometry Data and Tools to Transform Dysmorphology* (FB00000861); FB-TK0 [32], *3D White Light Photogrammetry Images of North American Children from Denver and San Francisco* (FB00000892); FB-VWP [33], *3D Facial Norms* (FB00000491.01); and FB-TX4 [34], *3D Facial Images—Tanzania* (FB00000667.01). Access to restricted data must be requested directly through FaceBase in accordance with its data access policies.

## Code Availability

The custom code for data preprocessing, model training, inference, and evaluation will be made publicly available upon publication at facebase 3d encoder benchmark. Scripts for cohort sensitivity analysis, repeated-scan aggregation, prototype-based diagnostic retrieval, and anatomical interpretability analysis will also be provided. Subject-level data partitions will be shared where permitted by FaceBase data-sharing requirements to facilitate reproducibility.

## Acknowledgements

This work was supported in part by the National Institutes of Health (NIH) under grants U54TR005652 and OT2OD037960, and by the Children’s Hospital of Philadelphia (CHOP) Research Institute. We acknowledge the FaceBase Consortium and the investigators who collected, curated, and contributed the 3D facial imaging datasets used in this study. We also thank the FaceBase Data Management Hub for facilitating access to these craniofacial research resources. FaceBase is supported by the National Institute of Dental and Craniofacial Research (NIDCR), part of the NIH. The content is solely the responsibility of the authors and does not necessarily represent the official views of the NIH.

## Supplementary Results

### Study overview and 3D analysis framework

Supplementary Fig. S1 provides additional details of the study design and analysis framework. Panel (a) illustrates how template fitting can reduce the measured deviation of facial morphology from a normative template, motivating the use of facial representations without template fitting. Panels (b–d) summarize the self-reported ancestry distributions, cohort sizes, and contributions of the six FaceBase cohorts to the benchmark tasks. Panel (e) illustrates the point-cloud encoding pipeline and the two inference strategies, softmax classification and prototype inference, at both scan and patient levels.

**Figure S1:**
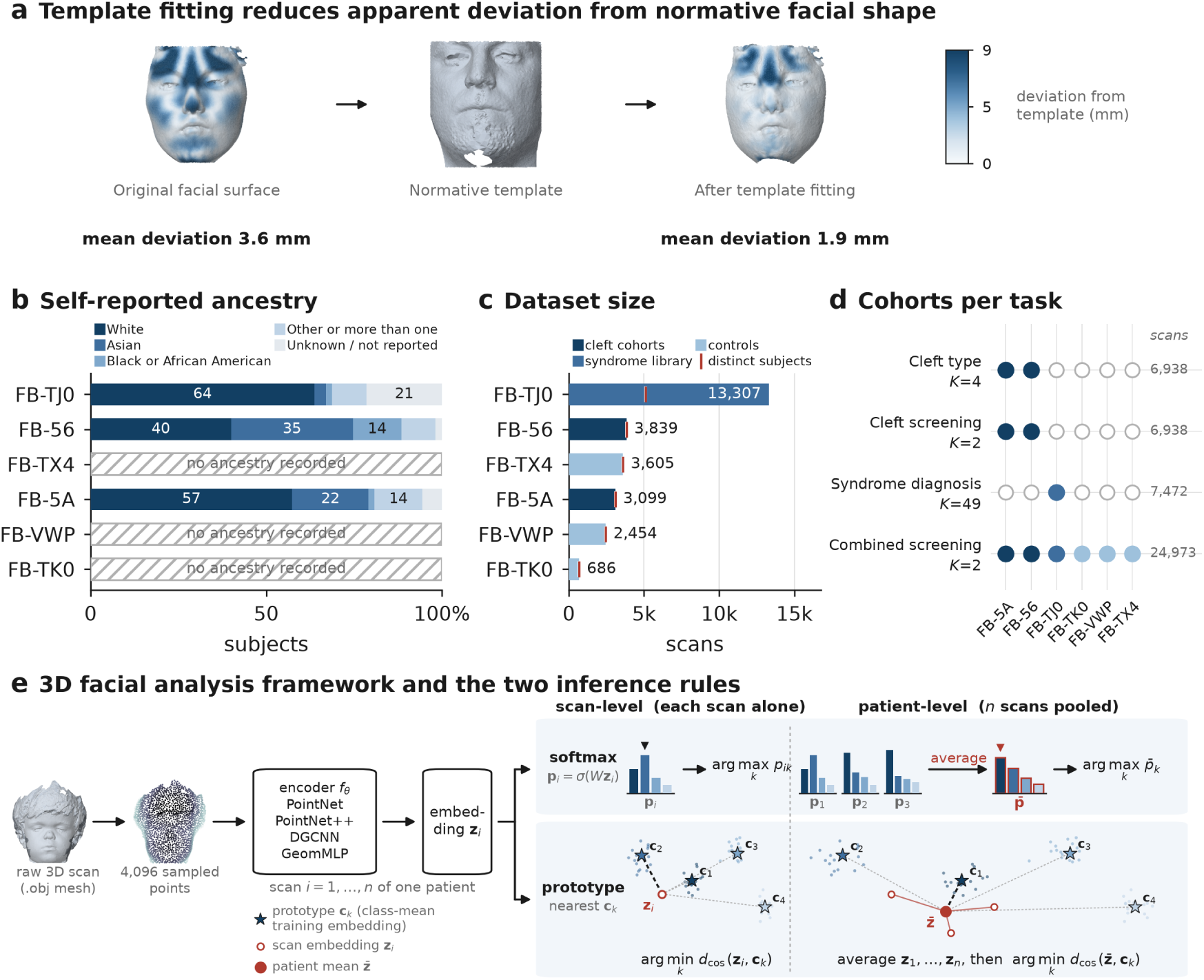
Study overview and 3D facial analysis framework. **(a)** Effect of template fitting on measured facial shape deviation. A representative 22q11.2 deletion syndrome scan is shown before fitting (left), alongside a normative control template (center) and after regularized non-rigid fitting (right). Colors indicate geometric deviation from the template, with mean deviation decreasing from 3.6 mm to 1.9 mm after fitting. This example illustrates how template fitting can reduce apparent morphological differences. **(b)** Distribution of self-reported ancestry across six FaceBase cohorts. Ancestry information was available for FB-TJ0, FB-56, and FB-5A but was not reported for FB-TX4, FB-VWP, or FB-TK0. **(c)** Numbers of processed scans (bars) and distinct subjects (red ticks) in each cohort. FB-TJ0 contains repeated acquisitions from many subjects. **(d)** Contributions of the six cohorts to the four benchmark tasks. Filled circles indicate cohort inclusion; *K* denotes the number of classes, and numbers at right indicate included scans. **(e)** 3D facial analysis framework. Each scan was represented by a 4,096-point cloud without template registration or dense correspondence and processed using one of four encoders: GeomMLP, PointNet, PointNet++, or DGCNN. At scan level, predictions were obtained independently for each acquisition. At patient level, softmax probabilities were averaged across scans, whereas prototype inference averaged scan embeddings before comparison with class prototypes.

### Cohort and demographic sensitivity of combined screening

Supplementary Table S1 provides detailed results for the cohort-specific, age-restricted, age- and sex-matched, and demographic-adjusted screening analyses. It reports the evaluation sample sizes, performance across independently trained PointNet++ models, and ensemble estimates with confidence intervals where available.

**Table S1:** Within-cohort and demographic sensitivity analyses of combined screening performance. All analyses used subject-level predictions from the existing combined-task Point-Net++ models without retraining. PointNet++ results are reported as mean *±* SD across five independently trained models, with five inference seeds averaged within each model. Ensemble AUCs were calculated by averaging predictions across all 25 model–inference-seed combinations; where reported, 95% confidence intervals were obtained from 2,000 subject-level bootstrap resamples. Sample sizes (*n*) indicate affected/unaffected test subjects, and ages are reported as medians in years. *^a^*Affected and unaffected subjects were matched 1:1 within the same cohort and sex, without replacement, with an absolute age difference *≤* 2 years. *^b^*Age-only logistic regression was trained on non-test subjects and evaluated on held-out test subjects. *^c^*Age- and sex-adjusted logistic regression was evaluated using stratified five-fold cross-validation repeated 20 times within the held-out test subjects, with or without the existing PointNet++ score.

| Evaluation | *n*<br>(aff./unaff.) | Median age, y<br>(aff./unaff.) | AUC<br>mean ± SD | Ensemble AUC<br>[95% CI] |
|---|---|---|---|---|
| *All cohorts combined* |  |  |  |  |
| PointNet++, all test subjects | 697 / 1,961 | – | 0.954 ± 0.004 | 0.967 [0.961, 0.973] |
| Cohort identity only | 697 / 1,961 | – | 0.912 | – |
| Age only<sup>*b*</sup> | 2,012 total | – | 0.743 | – |
| *FB-TJ0 (syndromic)* |  |  |  |  |
| PointNet++, all test subjects | 524 / 124 | 13.1 / 32.8 | 0.828 ± 0.017 | 0.861 [0.817, 0.898] |
| PointNet++, age/sex-matched<sup>*a*</sup> | 107 / 107 | 31.0 / 30.6 | 0.714 ± 0.023 | 0.743 [0.670, 0.808] |
| PointNet++, age <18 y | 350 / 22 | 8.8 / 11.3 | 0.606 ± 0.044 | 0.626 [0.489, 0.761] |
| Cohort identity only | 524 / 124 | – | 0.500 | – |
| Age only<sup>*b*</sup> | 635 total | – | 0.782 | – |
| Age + sex, CV<sup>*c*</sup> | 520 / 115 | – | 0.776 | – |
| Age + sex + PointNet++, CV<sup>*c*</sup> | 520 / 115 | – | 0.833 ± 0.012 | 0.854 |
| *Cleft cohorts (FB-5A + FB-56)* |  |  |  |  |
| PointNet++, all test subjects | 173 / 862 | 7.3 / 28.9 | 0.874 ± 0.014 | 0.902 [0.873, 0.926] |
| PointNet++, age/sex-matched<sup>*a*</sup> | 147 / 147 | 8.3 / 8.4 | 0.749 ± 0.023 | 0.780 [0.726, 0.832] |
| PointNet++, age <18 y | 150 / 247 | 6.0 / 9.0 | 0.817 ± 0.024 | 0.852 [0.810, 0.889] |
| Cohort identity only | 173 / 862 | – | 0.551 | – |
| Age only<sup>*b*</sup> | 1,035 total | – | 0.839 | – |
| Age + sex, CV<sup>*c*</sup> | 173 / 862 | – | 0.857 | – |
| Age + sex + PointNet++, CV<sup>*c*</sup> | 173 / 862 | – | 0.904 ± 0.005 | 0.918 |
| *FB-5A* |  |  |  |  |
| PointNet++, all test subjects | 96 / 390 | 9.0 / 29.0 | 0.861 ± 0.025 | 0.893 [0.854, 0.927] |
| PointNet++, age/sex-matched<sup>*a*</sup> | 95 / 95 | 9.0 / 9.0 | 0.786 ± 0.031 | 0.815 [0.752, 0.873] |
| PointNet++, age <18 y | 81 / 135 | 8.0 / 9.0 | 0.803 ± 0.034 | 0.837 [0.779, 0.887] |
| Age only<sup>*b*</sup> | 486 total | – | 0.785 | – |
| Age + sex, CV<sup>*c*</sup> | 96 / 390 | – | 0.802 | – |
| Age + sex + PointNet++, CV<sup>*c*</sup> | 96 / 390 | – | 0.887 ± 0.012 | 0.905 |
| *FB-56* |  |  |  |  |
| PointNet++, all test subjects | 77 / 472 | 5.2 / 28.9 | 0.882 ± 0.009 | 0.907 [0.867, 0.943] |
| PointNet++, age/sex-matched<sup>*a*</sup> | 54 / 54 | 8.1 / 8.0 | 0.696 ± 0.041 | 0.727 [0.621, 0.816] |
| PointNet++, age <18 y | 69 / 112 | 4.9 / 9.4 | 0.835 ± 0.020 | 0.868 [0.809, 0.921] |
| Age only<sup>*b*</sup> | 549 total | – | 0.885 | – |
| Age + sex, CV<sup>*c*</sup> | 77 / 472 | – | 0.899 | – |
| Age + sex + PointNet++, CV<sup>*c*</sup> | 77 / 472 | – | 0.916 ± 0.006 | 0.925 |

### Patient-level aggregation on the combined screening task

Patient-level aggregation provided little benefit for combined affected-versus-unaffected screening (Supplementary Table S2). Across the four models, macro-*F*_1_ decreased by 0.8 to 1.7 percentage points, with the paired bootstrap confidence interval excluding zero in all five training runs for PointNet, PointNet++ and DGCNN, and in one of five for GeomMLP. Changes in accuracy were small and were consistently resolved across all five retrainings for every model except DGCNN, while macro AUC changed by less than one percentage point for every model (*−*0.2 to +0.8).

Thus, unlike fine-grained syndrome diagnosis, broad screening did not gain additional discriminative performance from combining repeated scans. Where scan-level performance was already high, subject-level aggregation produced only small changes in accuracy and AUC while reducing class-balanced performance as measured by macro-*F*_1_. This contrast indicates that the benefit of repeated-scan aggregation is greatest for more difficult, fine-grained diagnostic tasks.

The aggregation analysis for screening was limited to the combined affected-versus-unaffected task. In the binary cleft screening cohort, each subject contributed a single scan, making scan-level and subject-level prediction identical by construction. The effect of repeated-scan aggregation could therefore not be evaluated independently in that cohort.

**Table S2:** Effect of patient-level aggregation on combined affected-versus-unaffected screening. Scan- and subject-level performance is shown for four facial-shape models using softmax inference on 3,639 held-out scans from 2,658 subjects. Subject-level predictions were obtained by averaging class-probability vectors across repeated scans from the same individual. Values are mean *±* standard deviation across five independently trained models using the same subject-level data split. For each model, Δ denotes the paired subject-minus-scan difference in percentage points. Acc, accuracy; macro-*F*_1_, macro-averaged *F*_1_; AUC, macro one-versus-rest area under the receiver operating characteristic curve.

| Model | Prediction level | Acc | Macro-*F*<sub>1</sub> | AUC |
|---|---|---|---|---|
| GeomMLP | Scan | 0.806 ± 0.004 | 0.800 ± 0.003 | 0.886 ± 0.003 |
|  | Subject | 0.837 ± 0.007 | 0.792 ± 0.005 | 0.894 ± 0.003 |
|  | Δ (pp) | +3.0 ± 0.4 | −0.8 ± 0.3 | +0.8 ± 0.1 |
| PointNet | Scan | 0.867 ± 0.010 | 0.864 ± 0.010 | 0.939 ± 0.004 |
|  | Subject | 0.881 ± 0.012 | 0.849 ± 0.012 | 0.939 ± 0.005 |
|  | Δ (pp) | +1.4 ± 0.3 | −1.5 ± 0.3 | +0.0 ± 0.1 |
| PointNet++ | Scan | 0.893 ± 0.007 | 0.890 ± 0.007 | 0.956 ± 0.002 |
|  | Subject | 0.900 ± 0.007 | 0.875 ± 0.008 | 0.954 ± 0.004 |
|  | Δ (pp) | +0.8 ± 0.1 | −1.5 ± 0.2 | −0.2 ± 0.2 |
| DGCNN | Scan | 0.874 ± 0.005 | 0.871 ± 0.006 | 0.948 ± 0.005 |
|  | Subject | 0.883 ± 0.003 | 0.854 ± 0.004 | 0.946 ± 0.005 |
|  | Δ (pp) | +0.9 ± 0.3 | −1.7 ± 0.2 | −0.2 ± 0.1 |

### Controlled same-patient aggregation: patient counts, increments and distributions

Of the 385 test patients in the 49-class syndrome-diagnosis task, 102 contributed one scan, while 283, 166, 101, and 51 contributed at least two, three, four, and five scans, respectively. These groups represented 48, 46, 42, and 28 diagnoses. Patients with a single scan contributed to the evaluation-unit comparison (Fig. 4b) but not to the controlled same-patient comparisons.

### Scan-count effects

Supplementary Table S3 reports the incremental effects of combining additional scans within fixed patient groups. Both top-1 accuracy and the rank of the true syndrome generally improved with increasing scan count under softmax and prototype inference, with the largest gains occurring when adding the second scan. All incremental improvements were positive across the five trained models, and the corresponding 95% bootstrap confidence intervals excluded zero. Among patients with at least five scans, combining five rather than one increased top-1 accuracy by 12.3 percentage points under softmax inference and 18.7 percentage points under prototype inference, while improving the mean true-syndrome rank by 2.92 and 3.34 positions, respectively.

### Patient-level variability

Figure 4f summarizes the average change in true-syndrome rank for each patient when a second scan was added. Under prototype inference, the correct syndrome moved higher for 254 patients and lower for 29, compared with 210 and 69 under softmax inference. These counts include small nonzero changes in average rank. Requiring a minimum improvement or deterioration of 0.5 or 1 rank position reduced the number of patients classified as changed (188/16 and 144/15 under prototype inference; 145/40 and 105/21 under softmax inference). Overall, prototype inference showed more frequent positive patient-level rank changes, although individual responses to aggregation remained heterogeneous.

**Table S3:**
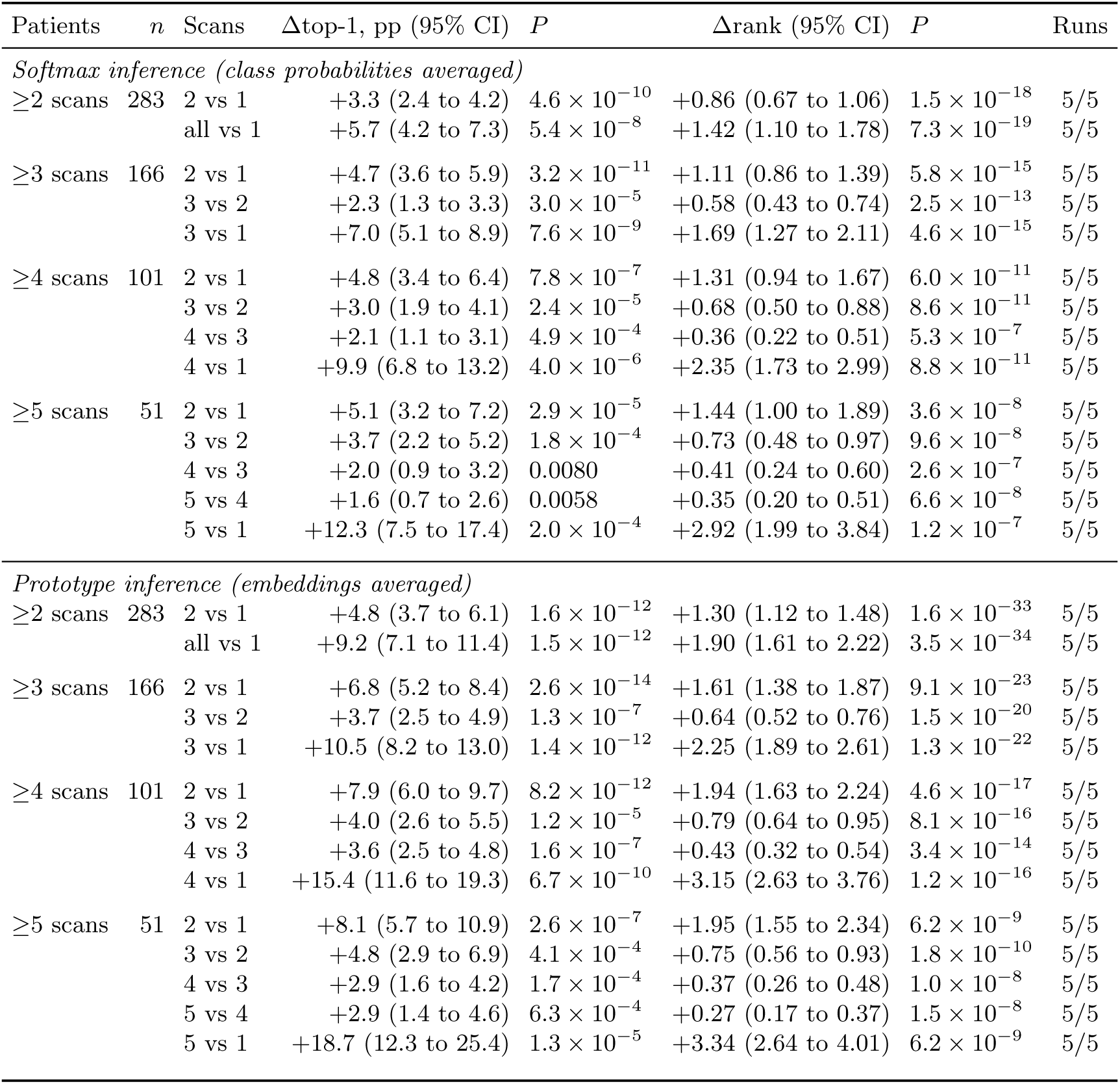
Controlled same-patient analysis of incremental scan aggregation. Results are based on five independently trained PointNet++ models for 49-class syndrome diagnosis, without retraining. Each block includes a fixed group of patients with at least the indicated number of scans. Comparisons evaluate additional scans within the same patients, averaging over all possible scan subsets or 300 randomly sampled subsets. Δtop-1 is the change in top-1 accuracy (percentage points), and Δrank is the improvement in the rank of the true syndrome among 49 candidates (positive values indicate better ranking). Values are means across five trained models, with inference seeds averaged within each model. Confidence intervals were obtained from 2,000 patient-level bootstrap resamples; *P* values were calculated using two-sided Wilcoxon signed-rank tests on paired patient-level differences. Runs indicates the number of trained models showing positive improvements in both metrics.

| Patients | *n* | Scans | Δtop-1, pp (95% CI) | *P* | Δrank (95% CI) | *P* | Runs |
|---|---|---|---|---|---|---|---|
| *Softmax inference (class probabilities averaged)* |  |  |  |  |  |  |  |
| ≥2 scans | 283 | 2 vs 1 | +3.3 (2.4 to 4.2) | 4.6 × 10<sup>−10</sup> | +0.86 (0.67 to 1.06) | 1.5 × 10<sup>−18</sup> | 5/5 |
|  |  | all vs 1 | +5.7 (4.2 to 7.3) | 5.4 × 10<sup>−8</sup> | +1.42 (1.10 to 1.78) | 7.3 × 10<sup>−19</sup> | 5/5 |
| ≥3 scans | 166 | 2 vs 1 | +4.7 (3.6 to 5.9) | 3.2 × 10<sup>−11</sup> | +1.11 (0.86 to 1.39) | 5.8 × 10<sup>−15</sup> | 5/5 |
|  |  | 3 vs 2 | +2.3 (1.3 to 3.3) | 3.0 × 10<sup>−5</sup> | +0.58 (0.43 to 0.74) | 2.5 × 10<sup>−13</sup> | 5/5 |
|  |  | 3 vs 1 | +7.0 (5.1 to 8.9) | 7.6 × 10<sup>−9</sup> | +1.69 (1.27 to 2.11) | 4.6 × 10<sup>−15</sup> | 5/5 |
| ≥4 scans | 101 | 2 vs 1 | +4.8 (3.4 to 6.4) | 7.8 × 10<sup>−7</sup> | +1.31 (0.94 to 1.67) | 6.0 × 10<sup>−11</sup> | 5/5 |
|  |  | 3 vs 2 | +3.0 (1.9 to 4.1) | 2.4 × 10<sup>−5</sup> | +0.68 (0.50 to 0.88) | 8.6 × 10<sup>−11</sup> | 5/5 |
|  |  | 4 vs 3 | +2.1 (1.1 to 3.1) | 4.9 × 10<sup>−4</sup> | +0.36 (0.22 to 0.51) | 5.3 × 10<sup>−7</sup> | 5/5 |
|  |  | 4 vs 1 | +9.9 (6.8 to 13.2) | 4.0 × 10<sup>−6</sup> | +2.35 (1.73 to 2.99) | 8.8 × 10<sup>−11</sup> | 5/5 |
| ≥5 scans | 51 | 2 vs 1 | +5.1 (3.2 to 7.2) | 2.9 × 10<sup>−5</sup> | +1.44 (1.00 to 1.89) | 3.6 × 10<sup>−8</sup> | 5/5 |
|  |  | 3 vs 2 | +3.7 (2.2 to 5.2) | 1.8 × 10<sup>−4</sup> | +0.73 (0.48 to 0.97) | 9.6 × 10<sup>−8</sup> | 5/5 |
|  |  | 4 vs 3 | +2.0 (0.9 to 3.2) | 0.0080 | +0.41 (0.24 to 0.60) | 2.6 × 10<sup>−7</sup> | 5/5 |
|  |  | 5 vs 4 | +1.6 (0.7 to 2.6) | 0.0058 | +0.35 (0.20 to 0.51) | 6.6 × 10<sup>−8</sup> | 5/5 |
|  |  | 5 vs 1 | +12.3 (7.5 to 17.4) | 2.0 × 10<sup>−4</sup> | +2.92 (1.99 to 3.84) | 1.2 × 10<sup>−7</sup> | 5/5 |
| *Prototype inference (embeddings averaged)* |  |  |  |  |  |  |  |
| ≥2 scans | 283 | 2 vs 1 | +4.8 (3.7 to 6.1) | 1.6 × 10<sup>−12</sup> | +1.30 (1.12 to 1.48) | 1.6 × 10<sup>−33</sup> | 5/5 |
|  |  | all vs 1 | +9.2 (7.1 to 11.4) | 1.5 × 10<sup>−12</sup> | +1.90 (1.61 to 2.22) | 3.5 × 10<sup>−34</sup> | 5/5 |
| ≥3 scans | 166 | 2 vs 1 | +6.8 (5.2 to 8.4) | 2.6 × 10<sup>−14</sup> | +1.61 (1.38 to 1.87) | 9.1 × 10<sup>−23</sup> | 5/5 |
|  |  | 3 vs 2 | +3.7 (2.5 to 4.9) | 1.3 × 10<sup>−7</sup> | +0.64 (0.52 to 0.76) | 1.5 × 10<sup>−20</sup> | 5/5 |
|  |  | 3 vs 1 | +10.5 (8.2 to 13.0) | 1.4 × 10<sup>−12</sup> | +2.25 (1.89 to 2.61) | 1.3 × 10<sup>−22</sup> | 5/5 |
| ≥4 scans | 101 | 2 vs 1 | +7.9 (6.0 to 9.7) | 8.2 × 10<sup>−12</sup> | +1.94 (1.63 to 2.24) | 4.6 × 10<sup>−17</sup> | 5/5 |
|  |  | 3 vs 2 | +4.0 (2.6 to 5.5) | 1.2 × 10<sup>−5</sup> | +0.79 (0.64 to 0.95) | 8.1 × 10<sup>−16</sup> | 5/5 |
|  |  | 4 vs 3 | +3.6 (2.5 to 4.8) | 1.6 × 10<sup>−7</sup> | +0.43 (0.32 to 0.54) | 3.4 × 10<sup>−14</sup> | 5/5 |
|  |  | 4 vs 1 | +15.4 (11.6 to 19.3) | 6.7 × 10<sup>−10</sup> | +3.15 (2.63 to 3.76) | 1.2 × 10<sup>−16</sup> | 5/5 |
| ≥5 scans | 51 | 2 vs 1 | +8.1 (5.7 to 10.9) | 2.6 × 10<sup>−7</sup> | +1.95 (1.55 to 2.34) | 6.2 × 10<sup>−9</sup> | 5/5 |
|  |  | 3 vs 2 | +4.8 (2.9 to 6.9) | 4.1 × 10<sup>−4</sup> | +0.75 (0.56 to 0.93) | 1.8 × 10<sup>−10</sup> | 5/5 |
|  |  | 4 vs 3 | +2.9 (1.6 to 4.2) | 1.7 × 10<sup>−4</sup> | +0.37 (0.26 to 0.48) | 1.0 × 10<sup>−8</sup> | 5/5 |
|  |  | 5 vs 4 | +2.9 (1.4 to 4.6) | 6.3 × 10<sup>−4</sup> | +0.27 (0.17 to 0.37) | 1.5 × 10<sup>−8</sup> | 5/5 |
|  |  | 5 vs 1 | +18.7 (12.3 to 25.4) | 1.3 × 10<sup>−5</sup> | +3.34 (2.64 to 4.01) | 6.2 × 10<sup>−9</sup> | 5/5 |

### Per-class effects of prototype inference

Section 2.5 shows that prototype inference produced improvements in accuracy and macro-*F*_1_ but little change in macro AUC on the 49-class syndrome-diagnosis task. Supplementary Figs. S2 and S3 examine the class-specific changes underlying this result.

Supplementary Fig. S2 shows the change in per-class *F*_1_ from softmax to prototype inference. Changes were calculated separately within each independently trained PointNet++ model and then averaged across the five training runs, with inference-seed variation averaged within each run. Using an absolute change of at most 0.01 as the threshold for a negligible difference, prototype inference increased *F*_1_ for 31 diagnoses, decreased it for 9, and produced only small changes for 8; an additional 1 diagnoses had *F*_1_ = 0 under both inference rules. The direction of change for individual diagnoses varied across retrainings, with individual models showing 21–26 improved and 11–20 reduced diagnoses. Thus, prototype inference redistributed class-specific performance, although the effect for any individual diagnosis was not necessarily consistent across training runs.

Supplementary Fig. S3 compares these changes in *F*_1_ with corresponding changes in one-versus-rest AUC. The mean absolute per-class change was 0.055 for *F*_1_ and 0.034 for AUC, indicating that prototype inference altered class-specific assignment performance more than class-wise discrimination (about 1.6-fold). Despite these class-specific shifts, macro AUC changed little overall (Section 2.5), indicating that improvements for some diagnoses were largely offset by reductions for others.

**Figure S2:**
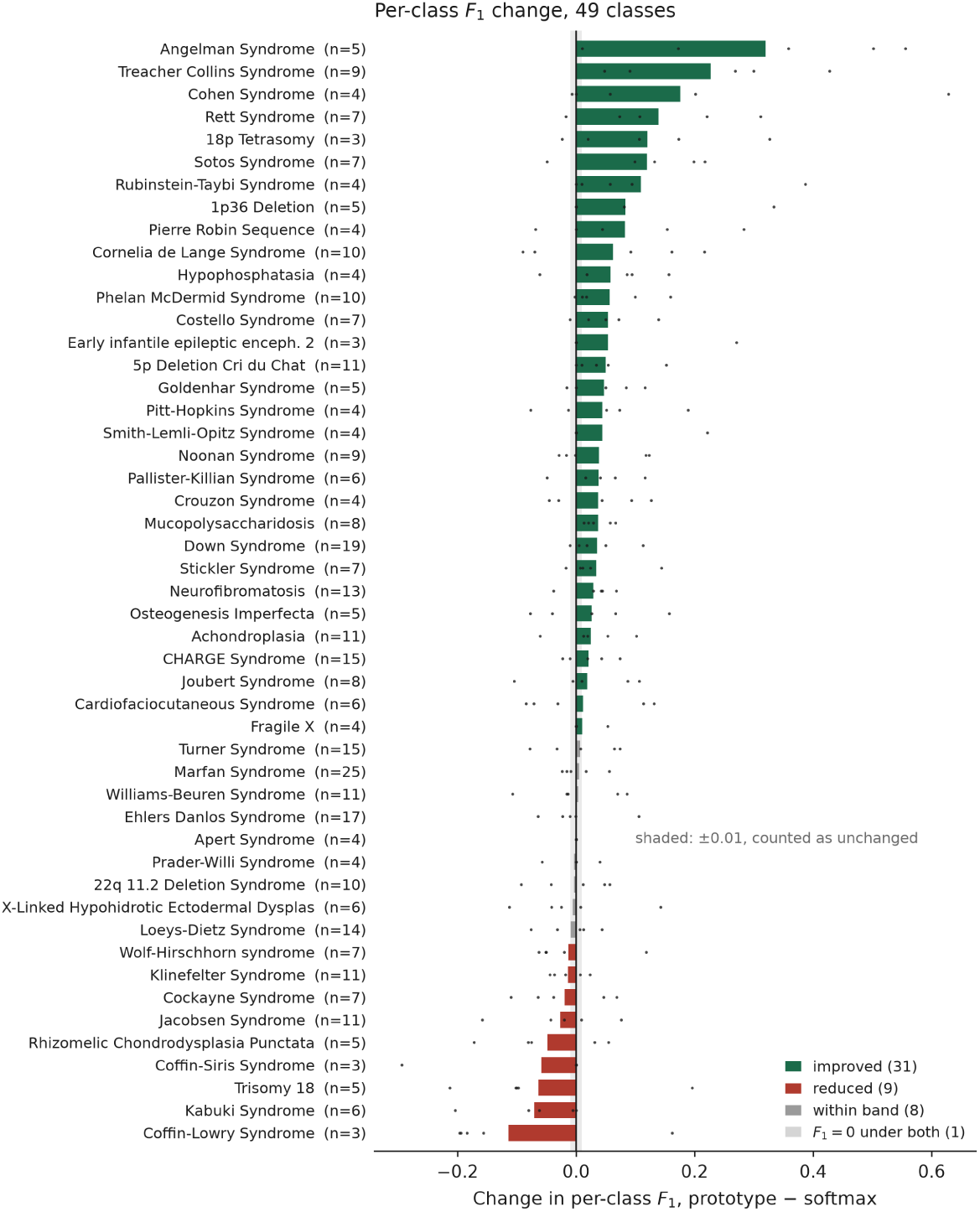
Class-specific changes in *F*_1_ under prototype inference for the 49-class syndrome-diagnosis task. Change in subject-level per-class *F*_1_ from softmax to prototype inference (prototype minus softmax) using PointNet++. Bars show the mean across five independently trained models, with five inference seeds averaged within each training run; dots show individual training runs. Each diagnosis is annotated with the number of test subjects in that class. Diagnoses are ordered by the direction and magnitude of the mean change. The shaded region denotes the *±*0.01 threshold used to classify changes as negligible.

**Figure S3:**
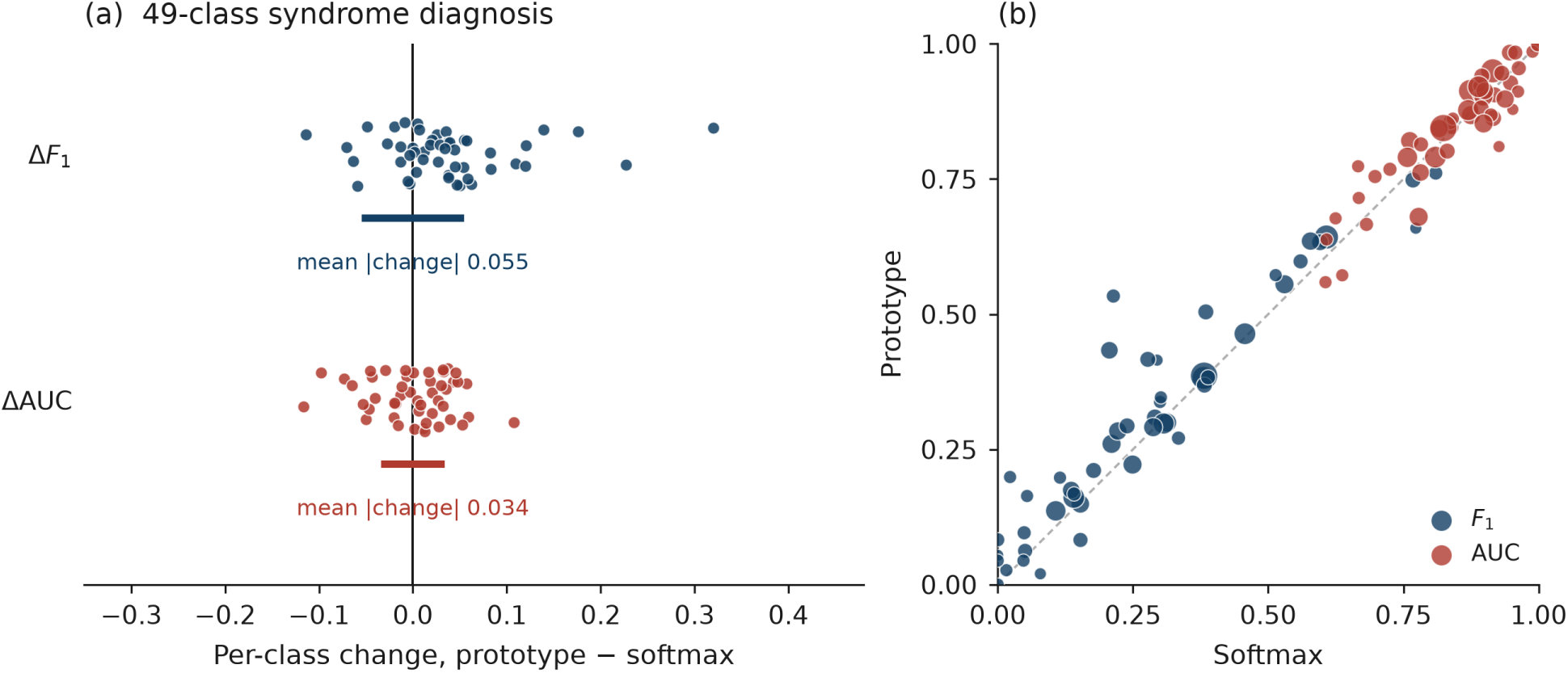
Prototype inference redistributes class-specific *F*_1_ and one-versus-rest AUC. Class-specific metrics were calculated within each of five independently trained PointNet++ models and then averaged across training runs, with five inference seeds averaged within each run. **(a)** Per-class changes from softmax to prototype inference for *F*_1_ and one-versus-rest AUC on the 49-class syndrome-diagnosis task. Each point represents one diagnosis, and horizontal bars indicate the mean absolute change across diagnoses. The same horizontal scale is used for both metrics. **(b)** Corresponding class-specific values under softmax and prototype inference. Marker area is proportional to the number of test subjects in each class, and the dashed diagonal denotes identical performance under the two inference rules.

### Sensitivity to the class-size threshold

The 49-class task retains diagnoses with at least 20 subjects. To assess the sensitivity of performance to this choice, we repeated training and evaluation at six additional thresholds, ranging from 10 subjects per diagnosis (55 diagnoses) to 75 subjects (12 diagnoses). PointNet++ subject-level prototype accuracy increased from 0.340 with 55 diagnoses to 0.477 with 12, whereas macro AUC varied more modestly, from 0.834 to 0.872 (Supplementary Fig. S4,b). PointNet++ and DGCNN outperformed PointNet and GeomMLP across all thresholds.

Because the test sets were nested, we further evaluated the same 173 subjects from the 12 diagnoses retained at the strictest threshold using models trained at every threshold (Supplementary Fig. S4). When prediction was restricted to these same 12 diagnoses, accuracy remained nearly constant across thresholds (PointNet++ 0.46–0.48; DGCNN 0.44–0.49). In contrast, when each model predicted among all diagnoses included during its training, PointNet++ accuracy on the same subjects decreased from 0.477 with 12 candidate diagnoses to 0.35–0.37 with 53–55. Thus, much of the lower top-1 accuracy at more permissive thresholds reflects the larger number of competing diagnoses rather than a substantial loss of recognition among the diagnoses shared across thresholds.

**Figure S4:**
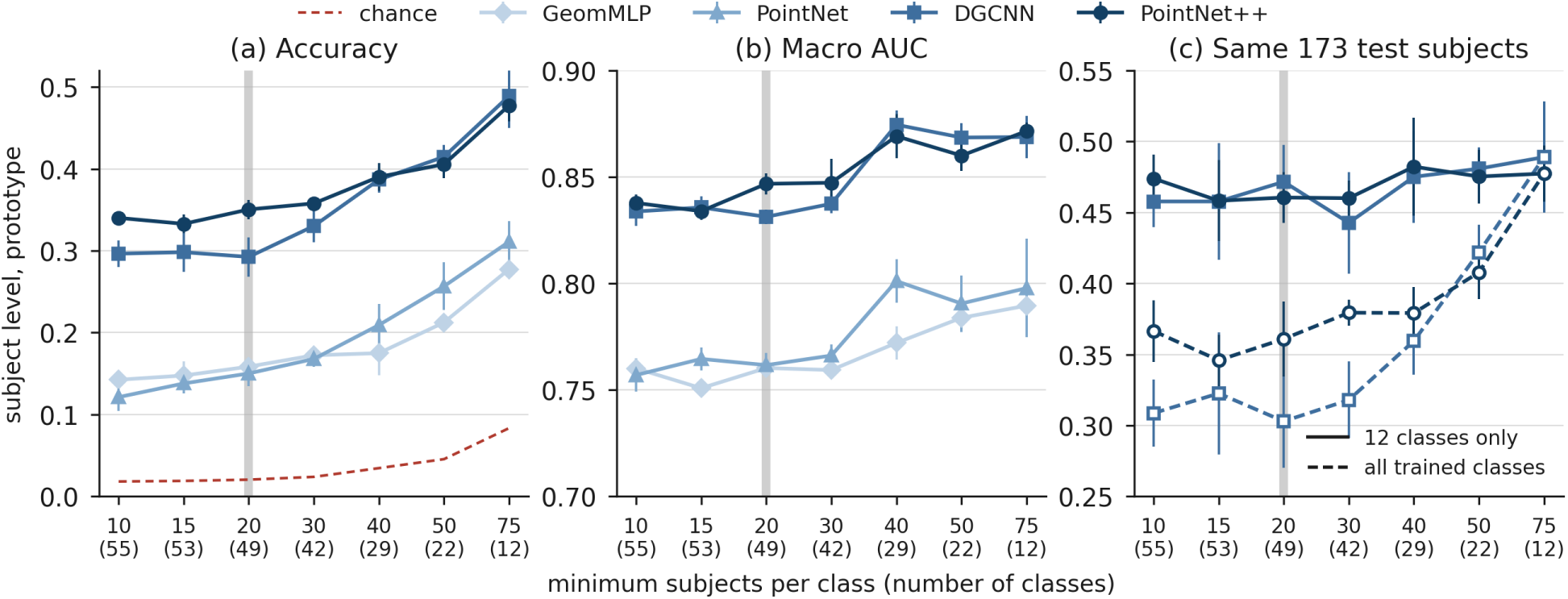
Sensitivity of syndrome diagnosis to the minimum class-size threshold. Diagnoses were retained if they had at least 50 scans and at least the indicated number of subjects; the number of retained diagnoses is shown in parentheses. The grey band marks the threshold used in the main analysis (20 subjects; 49 diagnoses). All thresholds use the same subject-level partition, resulting in nested test sets. Values show subject-level prototype inference as mean *±* standard deviation across five independently trained models. **(a)** Top-1 accuracy; dashed line indicates chance (1*/K*). **(b)** Macro one-versus-rest AUC. **(c)** Accuracy for the same 173 test subjects from the 12 diagnoses retained at the 75-subject threshold, evaluated using PointNet++ (circles) and DGCNN (squares) models trained at each threshold. Solid lines restrict prediction to the same 12 diagnoses, whereas dashed lines allow prediction among all diagnoses included at each training threshold.

### Structure of misclassification

The 49-class confusion matrix was examined to determine whether errors among phenotypically related diagnoses occurred more frequently than expected from their prevalence (Supplementary Fig. S5). We focused on Noonan syndrome, Costello syndrome, and cardiofaciocutaneous syndrome (CFC), the three RASopathies represented in the task.

Across five independently trained PointNet++ models, 17% of misclassified RASopathy subjects were assigned to another RASopathy. Because RASopathies accounted for 6% of the test set, within-family errors occurred about 2.9 times more often than expected from prevalence alone. The within-family fraction ranged from 6% to 33% across retrainings and exceeded the prevalence baseline in four of the five runs; in the fifth it matched it (5.9% versus 5.7%).

The most frequent within-family destination was Costello syndrome, which received 13% of RASopathy errors (63 of 469), mostly from CFC (Supplementary Fig. S5); the most frequent destination overall was neurofibromatosis (16%). Most RASopathy errors were nevertheless assigned to diagnoses outside the group (83%), most commonly neurofibromatosis, Joubert syndrome, and achondroplasia. Thus, the error pattern shows preferential confusion among related RASopathies, but within-family confusion still accounts for only a minority of their errors.

**Figure S5:**
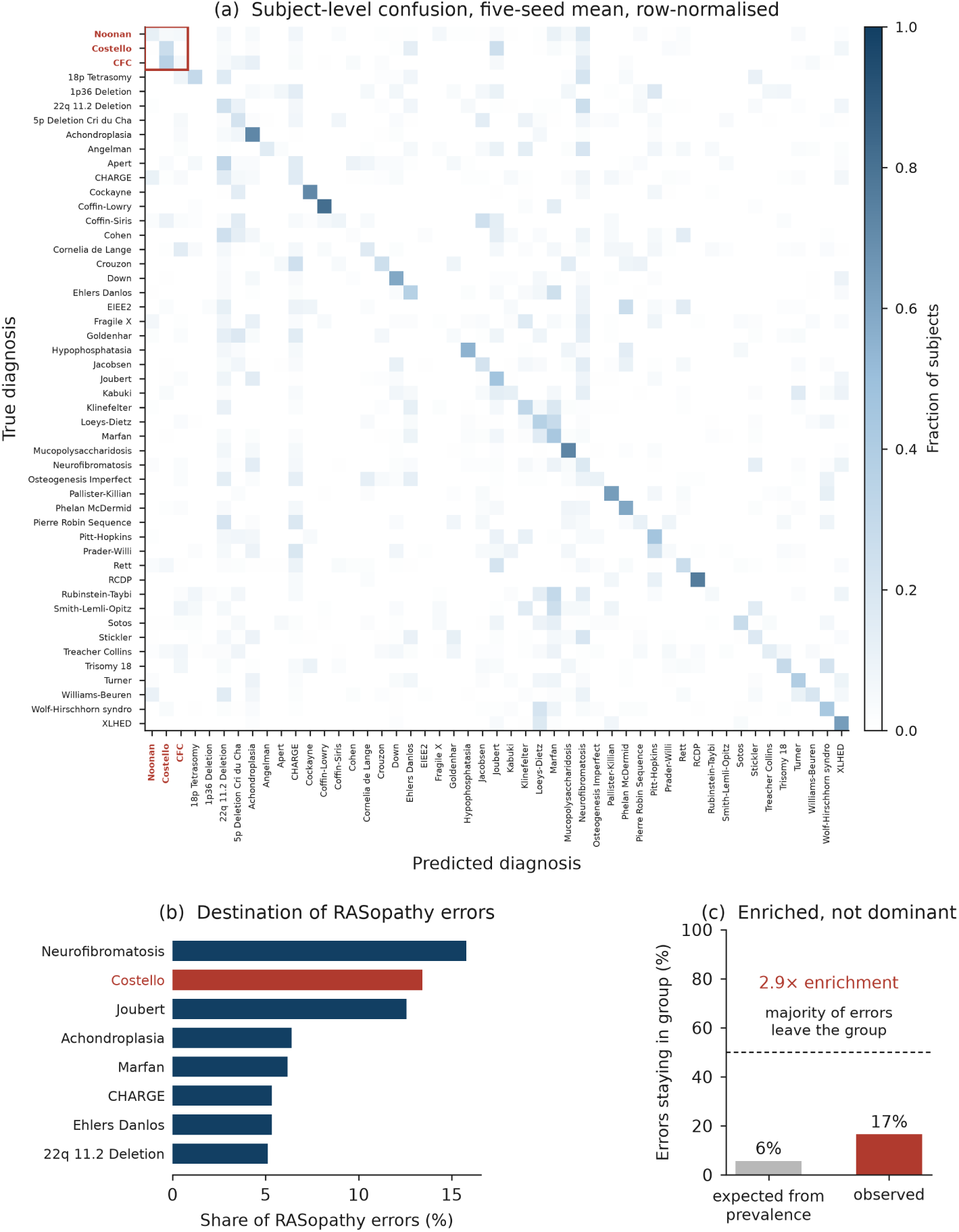
Structure of classification errors in the 49-class syndrome diagnosis task. **(a)** Row-normalized subject-level confusion matrix for PointNet++ with softmax inference, averaged across five independently trained models. Noonan syndrome, Costello syndrome, and cardiofaciocutaneous syndrome (CFC) are grouped and outlined in red. **(b)** Most frequent predicted diagnoses among misclassified RASopathy subjects; RASopathy destinations are shown in red. **(c)** Observed fraction of RASopathy errors assigned to another RASopathy compared with the fraction expected from their prevalence in the test set. Within-family errors were enriched relative to prevalence, although most RASopathy errors were assigned outside the group.

### Anatomical interpretability and retrieval characteristics across genetic syndromes

To further examine the learned facial representations, we analyzed nearest-neighbor retrieval and regional importance in correctly classified and misclassified subjects. For the five subjects shown in Fig. 7a, all three nearest training subjects shared the same diagnosis, supporting that the learned embeddings capture syndrome-associated facial similarities (Supplementary Fig. S6).

Misclassified subjects revealed several distinct patterns (Supplementary Fig. S7). First, confusion occurred between clinically related syndromes: cardiofaciocutaneous syndrome was consistently predicted as Costello syndrome, and Marfan syndrome as Loeys–Dietz syndrome, with all three nearest training subjects belonging to the respective predicted diagnoses. Second, a Down syndrome subject showed positive cheek and periorbital importance, similar to correctly classified examples, but was consistently misclassified as X-linked hypohidrotic ectodermal dysplasia. Finally, some misclassified subjects retrieved training subjects with the correct diagnosis among their nearest neighbors, as observed for Ehlers–Danlos and Williams–Beuren syndromes. These misclassifications highlight the challenges of differential diagnosis from 3D facial morphology, particularly when genetic syndromes share overlapping craniofacial phenotypes. Moreover, the presence of clinically relevant regional features or correctly labeled nearest neighbors does not necessarily translate into accurate syndrome discrimination.

**Figure S6:**
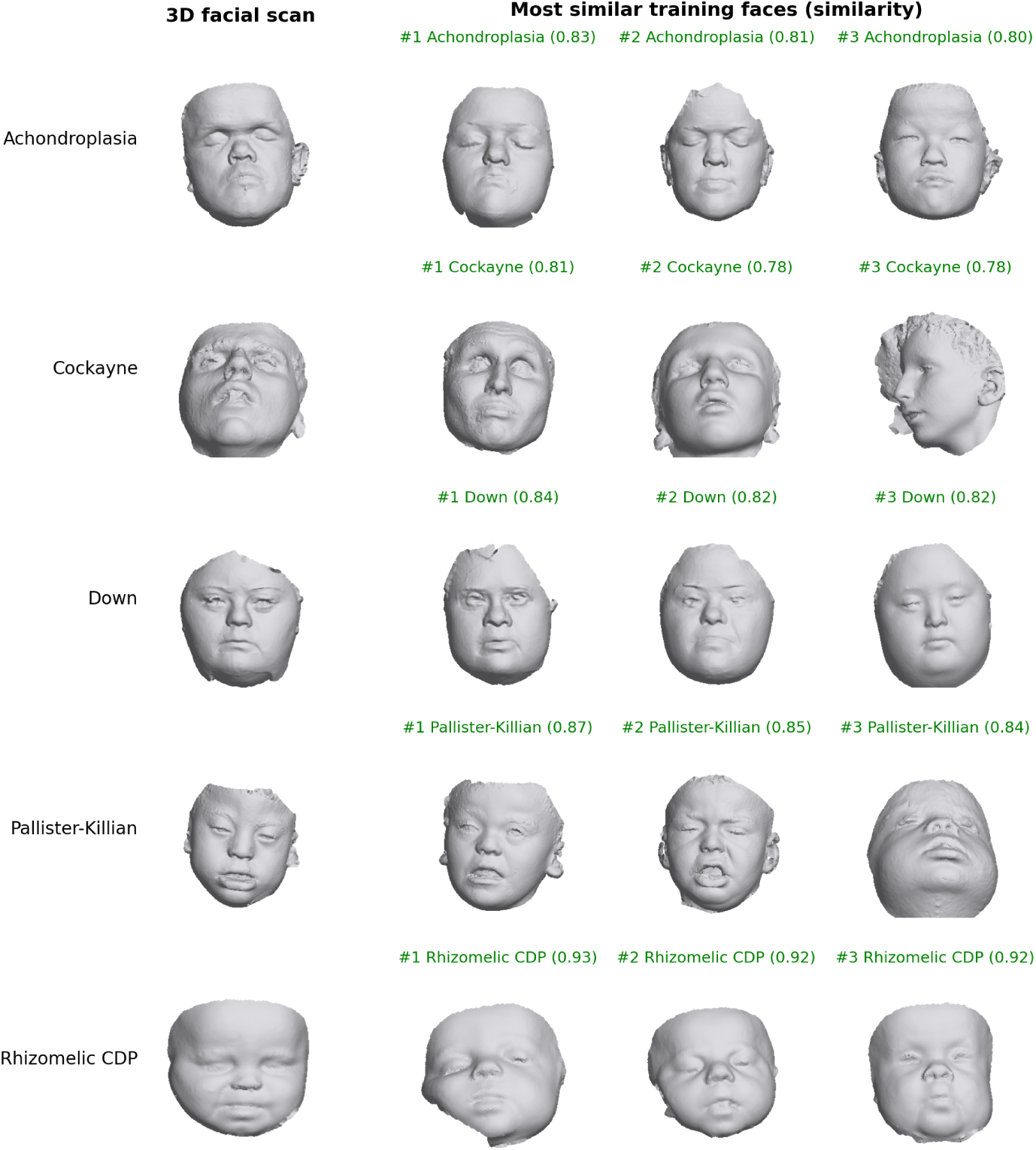
Nearest-neighbor retrieval for correctly classified genetic syndromes. For each subject shown in Fig. 7a, the three most similar training subjects are displayed, ranked by cosine similarity between subject-average embeddings averaged across five PointNet++ models. All 15 retrieved subjects share the diagnosis of the corresponding query subject (green labels), illustrating syndrome-associated similarity in the learned embedding space.

**Figure S7:**
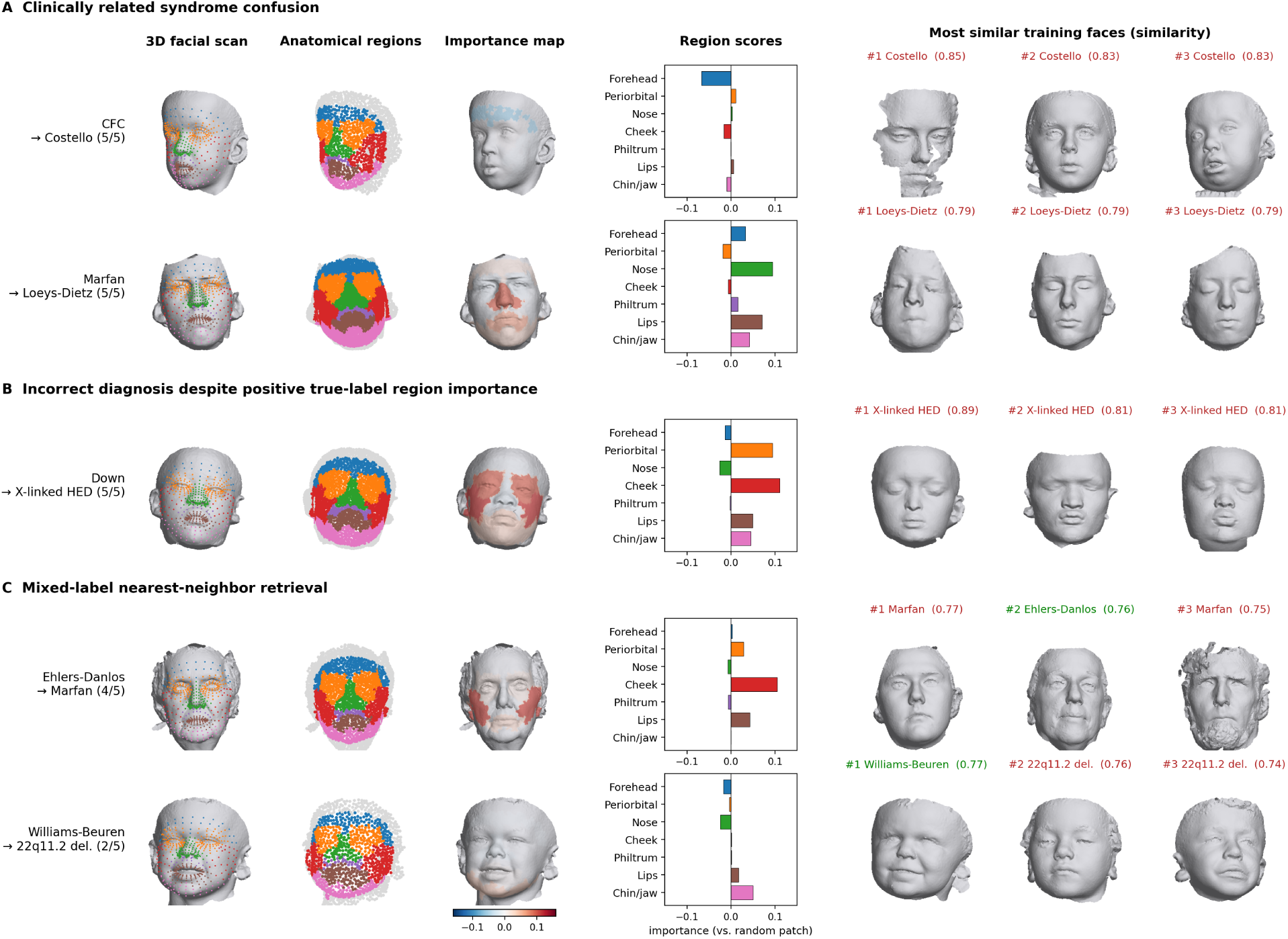
Anatomical region importance and nearest-neighbor retrieval in misclassified subjects. Test subjects misclassified by all five PointNet++ models are shown with their 3D facial scans, anatomical regions, importance maps, region scores, and three nearest training subjects. Importance scores were computed relative to the true diagnosis using the same region-occlusion procedure as in Fig. 7. Nearest neighbors were identified by cosine similarity of subject-average embeddings, averaged across the five models. Green labels indicate the correct diagnosis and red labels indicate a different diagnosis. Row labels show the true diagnosis and the most frequent incorrect prediction, with the number of models making that prediction. **(A)** Clinically related syndrome confusion: cardiofaciocutaneous syndrome (CFC) predicted as Costello syndrome and Marfan syndrome predicted as Loeys–Dietz syndrome, with all three nearest neighbors belonging to the predicted syndrome. **(B)** Incorrect diagnosis despite positive true-label importance: a Down syndrome subject showing positive cheek and periorbital contributions was predicted as X-linked hypohidrotic ectodermal dysplasia (X-linked HED). **(C)** Mixed-label nearest-neighbor retrieval: Ehlers–Danlos and Williams–Beuren subjects retrieved training subjects with the correct diagnosis despite being misclassified.

## References

[1] Judith E. Allanson, Christopher Cunniff, H. Eugene Hoyme, Julie McGaughran, Max Muenke, and Giovanni Neri. Elements of morphology: standard terminology for the head and face. American Journal of Medical Genetics Part A, 149A(1):6–28, 2009. doi: 10.1002/ajmg.a.32612.

[2] Peter Hammond. The use of 3D face shape modelling in dysmorphology. Archives of Disease in Childhood, 92(12):1120–1126, 2007. doi: 10.1136/adc.2006.103507.

[3] Peter A. Mossey, Julian Little, Ron G. Munger, Mike J. Dixon, and William C. Shaw. Cleft lip and palate. The Lancet, 374(9703):1773–1785, 2009. doi: 10.1016/S0140-6736(09)60695-4.

[4] Michael J. Dixon, Mary L. Marazita, Terri H. Beaty, and Jeffrey C. Murray. Cleft lip and palate: understanding genetic and environmental influences. Nature Reviews Genetics, 12(3): 167–178, 2011. doi: 10.1038/nrg2933.

[5] Tzung-Chien Hsieh and Peter M. Krawitz. Computational facial analysis for rare Mendelian disorders. American Journal of Medical Genetics Part C: Seminars in Medical Genetics, 193 (3):e32061, 2023. doi: 10.1002/ajmg.c.32061.

[6] Quentin Ferry, Julia Steinberg, Caleb Webber, David R. FitzPatrick, Chris P. Ponting, Andrew Zisserman, and Christoffer Nellåker. Diagnostically relevant facial gestalt information from ordinary photos. eLife, 3:e02020, 2014. doi: 10.7554/eLife.02020.

[7] Yaron Gurovich, Yair Hanani, Omri Bar, Guy Nadav, Nicole Fleischer, Dekel Gelbman, Lina Basel-Salmon, Peter M. Krawitz, Susanne B. Kamphausen, Martin Zenker, Lynne M. Bird, and Karen W. Gripp. Identifying facial phenotypes of genetic disorders using deep learning. Nature Medicine, 25(1):60–64, 2019. doi: 10.1038/s41591-018-0279-0.

[8] Tzung-Chien Hsieh, Martin A. Mensah, Jean T. Pantel, Dione Aguilar, Omri Bar, Allan Bayat, et al. PEDIA: prioritization of exome data by image analysis. Genetics in Medicine, 21(12): 2807–2814, 2019. doi: 10.1038/s41436-019-0566-2.

[9] Tzung-Chien Hsieh, Aviram Bar-Haim, Shahida Moosa, Nadja Ehmke, Karen W. Gripp, Jean Tori Pantel, Magdalena Danyel, Martin Atta Mensah, Denise Horn, Stanislav Rosnev, et al. Gestaltmatcher facilitates rare disease matching using facial phenotype descriptors. Nature Genetics, 54(3):349–357, 2022. doi: 10.1038/s41588-021-01010-x.

10. Da Wu, Jingye Yang, Cong Liu, Tzung-Chien Hsieh, Elaine Marchi, Justin Blair, Peter Krawitz, Chunhua Weng, Wendy Chung, Gholson J. Lyon, Ian D. Krantz, Jennifer M. Kalish, and Kai Wang. GestaltMML: Enhancing rare genetic disease diagnosis through multimodal machine learning combining facial images and clinical texts. arXiv preprint arXiv:2312.15320, 2024.

[11] Jordan J. Bannister, Matthias Wilms, J. David Aponte, David C. Katz, Ophir D. Klein, Francois P. Bernier, Richard A. Spritz, Benedikt Hallgrımsson, and Nils D. Forkert. Comparing 2d and 3d representations for face-based genetic syndrome diagnosis. European Journal of Human Genetics, 31:1010–1016, 2023. doi: 10.1038/s41431-023-01308-w.

[12] Peter Hammond, Tim J. Hutton, Judith E. Allanson, Bernard Buxton, Linda E. Campbell, Jill Clayton-Smith, Dian Donnai, Annette Karmiloff-Smith, Kay Metcalfe, Kieran C. Murphy, et al. Discriminating power of localized three-dimensional facial morphology. American Journal of Human Genetics, 77(6):999–1010, 2005. doi: 10.1086/498396.

[13] Benedikt Hallgŕımsson, J. David Aponte, David C. Katz, Jordan J. Bannister, Sheri L. Riccardi, Nick Mahasuwan, Brenda L. McInnes, Tracey M. Ferrara, Danika M. Lipman, et al. Automated syndrome diagnosis by three-dimensional facial imaging. Genetics in Medicine, 2020.

[14] Kristina Aldridge, Simeon A. Boyadjiev, George T. Capone, Valerie B. DeLeon, and Joan T. Richtsmeier. Precision and error of three-dimensional phenotypic measures acquired from 3dMD photogrammetric images. American Journal of Medical Genetics Part A, 138A(3):247–253, 2005. doi: 10.1002/ajmg.a.30959.

[15] Carrie L. Heike, Kristen Upson, Erik Stuhaug, and Seth M. Weinberg. 3D digital stereophotogrammetry: a practical guide to facial image acquisition. Head & Face Medicine, 6:18, 2010. doi: 10.1186/1746-160X-6-18.

[16] Pui Ki Hung, Junqi Liu, and Zhiyi Shan. 3D photogrammetry-driven craniofacial analysis in orthodontics: a scoping review of recent applications. Bioengineering, 12(11):1263, 2025. doi: 10.3390/bioengineering12111263.

[17] J. J. Bannister et al. An invertible statistical shape model for 3d facial morphology. IEEE Journal of Biomedical and Health Informatics, 2022.

[18] S. S. Mahdi et al. A multiscale analysis of 3d facial shape for syndrome delineation. IEEE Access, 2022.

[19] S. S. Mahdi et al. A 3d facial phenotype space for syndrome analysis. IEEE Access, 2024.

[20] Peter Hammond, Tim J. Hutton, Judith E. Allanson, Linda E. Campbell, Raoul C. M. Hennekam, Sean Holden, Michael A. Patton, Adam Shaw, I. Karen Temple, Matthew Trotter, Kieran C. Murphy, and Robin M. Winter. 3D analysis of facial morphology. American Journal of Medical Genetics Part A, 126A(4):339–348, 2004. doi: 10.1002/ajmg.a.20665.

[21] Julie D. White, Alejandra Ortega-Castrillon, Harold Matthews, Arslan A. Zaidi, Omid Ekrami, Jonatan Snyders, Yi Fan, Tony Penington, Stefan Van Dongen, Mark D. Shriver, and Peter Claes. MeshMonk: Open-source large-scale intensive 3D phenotyping. Scientific Reports, 9(1): 6085, 2019. doi: 10.1038/s41598-019-42533-y.

[22] James Booth, Anastasios Roussos, Allan Ponniah, David Dunaway, and Stefanos Zafeiriou. Large scale 3D morphable models. International Journal of Computer Vision, 126(2–4):233–254, 2018. doi: 10.1007/s11263-017-1009-7.

[23] Jordan J. Bannister, Sebastian R. Crites, J. David Aponte, David C. Katz, Matthias Wilms, Ophir D. Klein, Francois P. J. Bernier, Richard A. Spritz, Benedikt Hallgrımsson, and Nils D. Forkert. Fully automatic landmarking of syndromic 3D facial surface scans using 2D images. Sensors, 20(11):3171, 2020. doi: 10.3390/s20113171.

[24] Hanne Hoskens, Dongjing Liu, Sahin Naqvi, Myoung Keun Lee, Ryan J Eller, Karlijne Indencleef, Julie D White, Jiarui Li, Maarten HD Larmuseau, Greet Hens, et al. 3d facial phenotyping by biometric sibling matching used in contemporary genomic methodologies. PLoS genetics, 17(5): e1009528, 2021.

[25] Charles R. Qi, Hao Su, Kaichun Mo, and Leonidas J. Guibas. Pointnet: Deep learning on point sets for 3d classification and segmentation. In IEEE Conference on Computer Vision and Pattern Recognition (CVPR), 2017.

[26] Charles R. Qi, Li Yi, Hao Su, and Leonidas J. Guibas. Pointnet++: Deep hierarchical feature learning on point sets in a metric space. In Advances in Neural Information Processing Systems (NeurIPS), 2017.

[27] Yue Wang, Yongbin Sun, Ziwei Liu, Sanjay E. Sarma, Michael M. Bronstein, and Justin M. Solomon. Dynamic graph cnn for learning on point clouds. ACM Transactions on Graphics (TOG), 38(5):1–12, 2019.

[28] RE Schuler, A Bugacov, JG Hacia, TV Ho, J Iwata, L Pearlman, BD Samuels, C Williams, Z Zhao, C Kesselman, et al. Facebase: a community-driven hub for data-intensive research. Journal of Dental Research, 101(11):1289–1298, 2022.

[29] Mary L. Marazita, Seth Weinberg, and Joel Anderton. Pittsburgh oral facial cleft studies: Extending the phenotype of nonsyndromic orofacial clefting (ofc1). FaceBase Consortium, 2025. URL 10.25550/5A-FJBJ.

[30] Mary L. Marazita, Seth Weinberg, and Joel Anderton. Oral-facial cleft families: Phenotype and genetics (ofc2). FaceBase Consortium, 2025. URL 10.25550/56-ES6P.

[31] Benedikt Hallgrimsson, Richard A. Spritz, Ophir Klein, and Washington Mio. Developing 3d craniofacial morphometry data and tools to transform dysmorphology. FaceBase Consortium, 2017. URL 10.25550/TJ0.

[32] Richard A. Spritz, Benedikt Hallgrimsson, and Ophir Klein. 3d white light photogrammetry images of north american children from denver and san francisco.. FaceBase Consortium, 2017. URL 10.25550/TK0.

[33] Seth Weinberg and Zachary Marazita, Mary L. and Raffensperger. 3d facial norms. FaceBase Consortium, 2015. URL 10.25550/VWP.

[34] Benedikt Hallgrimsson, Richard A. Spritz, and Ophir Klein. 3d facial images - tanzania. FaceBase Consortium, 2015. URL 10.25550/TX4.

[35] Jordan J. Bannister, Matthias Wilms, J. David Aponte, David C. Katz, Ophir D. Klein, Francois P. J. Bernier, Richard A. Spritz, Benedikt Hallgrımsson, and Nils D. Forkert. Detecting 3D syndromic faces as outliers using unsupervised normalizing flow models. Artificial Intelligence in Medicine, 134:102425, 2022. doi: 10.1016/j.artmed.2022.102425.

[36] Yury Kartynnik, Artsiom Ablavatski, Ivan Grishchenko, and Matthias Grundmann. Real-time facial surface geometry from monocular video on mobile GPUs. arXiv preprint arXiv:1907.06724, 2019.

